# USP9X deubiquitinase couples the pluripotency network and cell metabolism to regulate ESC differentiation potential

**DOI:** 10.1101/2020.01.13.904904

**Authors:** Maud de Dieuleveult, Marjorie Leduc, Eralda Salataj, Céline Ransy, Julien Dairou, Kengo Homma, Morgane Le Gall, Pascale Bossard, Anne Lombès, Frédéric Bouillaud, Laure Ferry, Pierre-Antoine Defossez, Jean-Charles Cadoret, Hidenori Ichijo, Stephen A. Wood, François Guillonneau, Ralf Dressel, Benoit Miotto

## Abstract

Embryonic stem cells (ESC) have the unique ability to differentiate into all three germ cell layers. ESC transition through different states of pluripotency in response to growth factor signals and environmental cues before becoming terminally differentiated. Here, we demonstrated, by a multi-omic strategy, that the deubiquitinase USP9X regulates the developmental potential of ESC, and their transition from a naive to a more developmentally advance, or primed, state of pluripotency. We show that USP9X facilitates developmental gene expression and induces modifications of the mitochondrial bioenergetics, including decreased routing of pyruvate towards its oxidation and reduced respiration. In addition, USP9X binds to the pluripotency factor ESRRB, regulates its abundance and the transcriptional levels of a subset of its target genes. Finally, under permissive culture conditions, depletion of Usp9X accelerates cell differentiation in all cell lineages. We thus identified a new regulator of naive pluripotency and show that USP9X couples ESRRB pluripotency transcriptional network and cellular metabolism, both of which are important for ESC fate and pluripotency.

## INTRODUCTION

Pluripotency is defined by the ability to self-renew and, under appropriate conditions, differentiate into all somatic cell lineages [1]. Embryonic stem cells (ESC) transition to different states of pluripotency, that can be recapitulated *in vitro* by culture conditions, and that correlates with different behaviours and abilities to contribute to blastocyst chimeras [1–3]. The dynamic nature of ESC relies on discrete changes in signalling pathways, gene expression and epigenetic regulations. It also depends on cellular metabolism and growth factor dependencies.

System biology as well as detailed genomic analyses have shown that these different states of pluripotency mainly relied on the expression of specific sets of transcription factors [1,2,4–8]. As such, transcription factors OCT4, ZFP281, ZIC3, SOX2 and SALL4 contribute to a general program to maintain pluripotency while additional factors act with them to reinforce this network or, on the contrary, prime ESC to specific cell lineages [3]. For instance, high levels of KLF4, KLF5, NANOG, REX1, ZFP42 and ESRRB favour naive pluripotency [2,7,9–11]. This state of pluripotency is less permissive to transcriptional and functional heterogeneity than the state of ESC maintained in leukemia inhibitory factor (LIF) and serum [12, 13]. Experimentally, ESC can be maintained in the naive state in stem-cell media that includes two kinase inhibitors, known as the “2i” condition, to block MAPK Erk and glycogen synthase kinase-3 [12]. On the contrary, transcription factor LEFTY2, OTX2 and POU3F1 are transiently expressed when ESC are committed to differentiation, a state define as “formative” or “poised” pluripotency [1,2,8,14,15].

The targets of this network affect different cellular features of ESC, including their interaction with the microenvironment, their shape and their cellular metabolism. In ESC compared to differentiated cells, oxidative phosphorylation (OXPHOS) is low and ATP synthesis is more dependent on glycolysis [16–20]. In addition, inhibition of pyruvate dehydrogenase (PDH) activity causes a significant increase in lactate levels and acidification of the microenvironment; while mTOR and HIF pathway activities favour glycolysis [21, 22]. The preference for glycolytic metabolism is further reinforced in ESC cultured in “2i” compared to ESC cultured in LIF + serum [18, 21]. These changes in cell metabolism or microenvironment composition accompanying changes in pluripotency status might actually play a pivotal role in the regulation of cell fate and commitment. The coordination of the activity of the pluripotency network with other unique cellular features of ESC, including cell metabolism, remains to be clarified [17,21,23].

Ubiquitination/deubiquitination plays a central role in the plasticity of stem cells by regulating the abundance of the main pluripotency transcription factors, chromatin organization, cell metabolism and the architecture of the cytoskeleton [24]. Depletion of E3 ubiquitin ligases and deubiquitinases causes specific alterations in the ESC proteome with variable consequences on pluripotency and ESC fate [25, 26]. For example, depletion of the deubiquitinase USP21 causes NANOG degradation which triggers ESC differentiation [27]. WWP2 promotes SOX2 and OCT4 degradation and regulates the balance between self-renewal and cell differentiation [28, 29]. Finally, inactivation of TRIM32, which regulates the stability of the pluripotency factors MYC and OCT4, is not required for the maintenance of pluripotency, but it facilitates somatic cell reprogramming into induced pluripotent cells [30]. Thus, deubiquitinases and E3 Ubiquitin ligases play diverse roles in regulating ESC transitions into different pluripotency stages.

Deubiquitinase USP9X (also known as FAM or Fat Facets in Mammals) is indispensable for embryonic development [31, 32]. It plays a critical role in tissue development and homeostasis as exemplified by its implication in developmental disorders and cancers [33–39]. USP9X is very highly expressed in several stem cell populations compared to their differentiated counterparts, as well as in induced pluripotent cells, thus indicating USP9X expression as a marker of stemness [4,40,41]. It associates with the stem cell transcription factors SALL4 and SOX2 in co-immunoprecipitation assays and it was identified in a couple of high-throughput genetic screens aimed at identifying regulators of murine ESC biology and essential genes in human haploid ESC [26,42–44]. However, despite compelling evidences suggesting that USP9X regulates ESC biology, its function has not been properly studied in ESC.

The studies described in this report investigate the role of Usp9X in self-renewal and early differentiation of ESC. Experiments were conducted in Usp9X-knock-down and Usp9X-mutated murine ESC. We show that depletion of USP9X alters the balance of pluripotency transcription factor expression and accelerates the differentiation of ESC under permissive growth conditions. At the molecular level, USP9X reinforces the pluripotency status of ESC through its action on specific key stem cell transcription factors and cell metabolism.

## RESULTS

### USP9X controls the differentiation potential of ESC

To obtain insight into the role of USP9X in murine ESC, we utilized ESC carrying a deletion of exon 2 of Usp9X generated in a previous study [45]. PCR and RT-qPCR analyses confirmed the deletion of exon 2 in Usp9X^E2/Y^ ESC (Supplementary Figure S1A). Analyses by western blot, immunofluorescence and flow cytometry further confirmed residual amounts of USP9X protein detectable in Usp9X^E2/Y^ ESC (Figure 1A and Supplementary Figure S1B-C).

**Figure 1:**
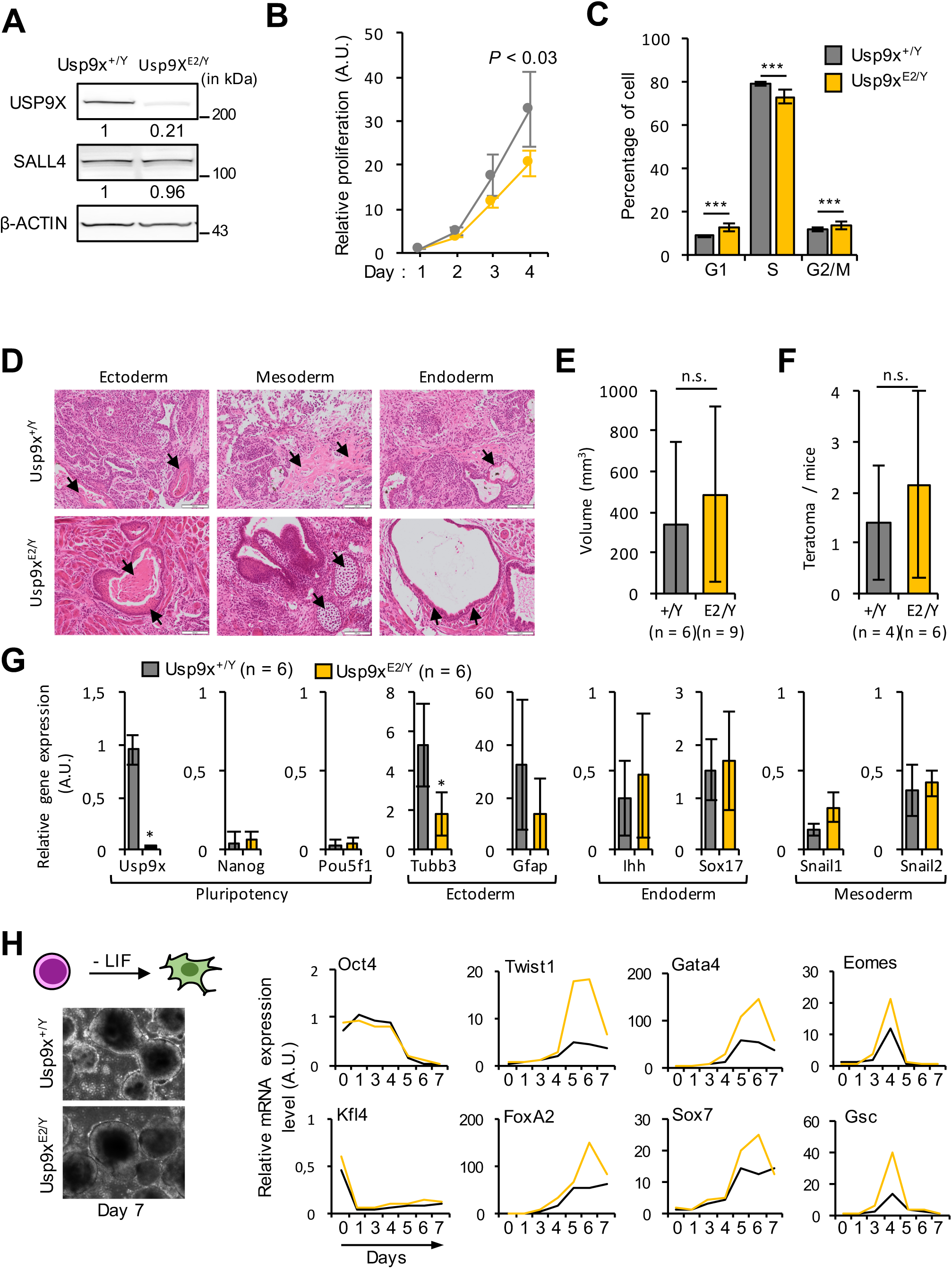
Molecular and phenotypical characterization of USP9X^E2/Y^ ESC. (**A**) Western blot analysis of USP9X, SALL4 and β-ACTIN levels in total protein extracts from USP9X^E2/Y^ and control ESC cultured in LIF + serum medium. Relative amounts of USP9X and SALL4 proteins normalised to β-ACTIN are indicated. (**B**) Proliferation of USP9X^E2/Y^ and control ESC cultured in LIF + serum medium and plated on gelatine (n = 3). *P*-value assessed by Wilcoxon signed rank test at day 4. (**C**) Cell cycle distribution of USP9X^E2/Y^ and control ESC cultured in LIF + serum medium. Ten thousand cells were analysed per condition (n = 3). ***, *P* < 0.00001 (assessed by Mann-Whitney test vs control). (**D**) Representative tissues seen in experimental teratomas derived from USP9X^E2/Y^ and control ESC. Tissues representing the three germ layers are indicated by arrows. Scale bar: 10 μm. (**E**) Quantification of the volume of teratomas (mean ± SD) derived from USP9X^E2/Y^ and control ESC. n.s., not significant (*P*-value assessed by Student t-test between groups). (**F**) Mean number of teratomas (± SD) derived from USP9X^E2/Y^ and control ESC per injected mice. n.s., not significant (*P*-value assessed by Student t-test between groups). (**G**) qRT-PCR analysis of pluripotency and cell differentiation markers in teratomas derived from USP9X^E2/Y^ and control ESC. Expression of pluripotency (Usp9X, Pou5f1 and Nanog), ectoderm (Gfap and Tubb3), endoderm (Ihh and Sox17) and mesoderm (Snail1 and Snail2) markers are presented as relative expression levels compared to Ppia. n.s., not significant; *, *P* < 0.05 (assessed by Student t-test between groups). (**H**) Usp9X regulates ESC differentiation upon LIF removal. Left panel: representative embryonic bodies from USP9X^E2/Y^ and control ESC seen at day 7 of differentiation. ESC differentiation was triggered by removal of LIF at day 0 and adding retinoic acid (5µM) at day 4 and 6. Right panel: RT-PCR analysis of pluripotency (Oct4 and Klf4) and differentiation (Twist1, FoxA2, Gata4, Sox7, Eomes and Gsc) markers expression levels in USP9X^E2/Y^ and control ESC at different days after differentiation.

Usp9X^E2/Y^ ESC proliferate slower than control isogenic cells in LIF + serum containing media (Figure 1B) and exhibit a slight accumulation in G1 and G2/M-phases of the cell cycle (Figure 1C). Nonetheless, Usp9X^E2/Y^ ESC express classical core pluripotency factors at similar levels to control ESC and do not aberrantly express the main differentiation markers of endoderm, mesoderm and ectoderm lineages (Supplementary Figure S1D). Importantly, when injected in immunodeficient mice, Usp9X^E2/Y^ ESC form teratomas containing tissues from all three germ layers, as their isogenic Usp9X^WT^ counterparts, indicating pluripotency *in vitro* (Figure 1D-F). RT-PCR analysis further confirmed that teratomas are formed from Usp9X^E2/Y^ ESC and show similar expression levels of the marker genes representing the three germ layers to control cells (Figure 1G). Thus, deletion of Usp9X exon 2 does not dramatically alter the maintenance of ESC or their ability to differentiate in all three germ cell layers upon injection in immune-deficient mice.

We next compared the kinetic of differentiation of Usp9X^E2/Y^ and control ESC *in vitro* by monitoring RNA levels of differentiation markers. We differentiated ESC in embryonic bodies (EB) by growing them in medium without LIF for 8 days in non-adherent dishes [46]. We confirmed the proper differentiation of control ESC by monitoring levels of pluripotency and differentiation markers from day 0 to day 8 (Supplementary Figure S1E). We observed that Usp9X^E2/Y^ ESC differentiate more efficiently compared to their isogenic counterpart (Figure 1H). This is clearly manifest by the early appearance of differentiation markers, such as Twist1, Gsc, Eomes, Gata4 and FoxoA2, and the higher amplitude of their induction (Figure 1H). The down-regulation of pluripotency markers occurs with similar kinetics in Usp9X^E2/Y^ and control ESC, and is concomitant with USP9X down-regulation (Supplementary Figure S1E). We further addressed the role of USP9X by studying the differentiation of Usp9X^E2/Y^ ESC in extra-embryonic endoderm (XEN) stem cells (Supplementary Figure S1F) [47]. Again, the differentiation of Usp9X^E2/Y^ was more efficient than their control counterparts, as illustrated by the higher levels of induction of FoxA2, Sox17 and Sox7 (Supplementary Figure S1G). Of note, USP9X was still expressed 8 days after starting XEN differentiation (Supplementary Figure S1F). These data indicate that the deletion of Usp9X exon 2 potentiates cell differentiation in different cell lineages independently of Usp9X mRNA kinetics during differentiation.

To further validate that it is the depletion of Usp9X in USP9X^E2/Y^ ESC that affects ESC features, we designed three short hairpin RNA (shRNA) constructs targeting different regions of Usp9X transcript (Usp9X^KD^). Knockdown efficiency was confirmed by western blot and label-free mass-spectrometry analysis (Supplementary Figure S2A-B). Usp9X^KD^ ESC did not exhibit obvious changes in the DNA replication program, cell proliferation and cell morphology compared to control cells (Supplementary Figure S2C-D). In addition, they retained alkaline phosphatase staining, a marker of pluripotency (Supplementary Figure S2E). Upon removal of LIF, expression of differentiation markers Gata4, Eomes, Foxa2 and Sox7 was induced faster and with higher amplitude in Usp9X^KD^ ESC compared to control cells, while expression of pluripotency factors Oct4 and Klf4 was down-regulated with very similar kinetics as in control shRNA transfected cells (Supplementary Figure S2F). These experiments showed that depletion of USP9X recapitulates the phenotype of USP9X^E2/Y^ cells.

Taken together, the results demonstrate that Usp9X regulates the regulatory network controlling ESC identity, and that its depletion accelerates ESC differentiation under permissive culture conditions.

### USP9X regulates glycolysis and naive pluripotency factor abundance

To understand the underlying mechanisms of USP9X depletion on ESC biology, we carried out an analysis of the proteome of ESC 64 hours after electroporation of USP9X or control short-hairpin RNAs (i.e. to allow 48h of puromycin selection post electroporation). We choose this approach to evaluate the response of the proteome to acute depletion of Usp9X, rather than in the adapted Usp9X^E2/Y^ mutant background. Among >6000 proteins detected, we identified 248 proteins in Usp9X^KD^ ESC with more than 1.1 fold change abundance and *P*-value < 0.05 as compared to control (Figure 2A; Supplementary Table 1). We validated by western blot several USP9X potential targets (Figure 2B). Among these 248 modified proteins, many were previously identified as potential targets of E3 ubiquitin ligases, as substrates of proteasome or as readers of ubiquitin-linkages in (human and mouse) ESC, indicating a potential regulation by Usp9X enzymatic activity (Supplementary Table 2) [25,26,48].

**Figure 2:**
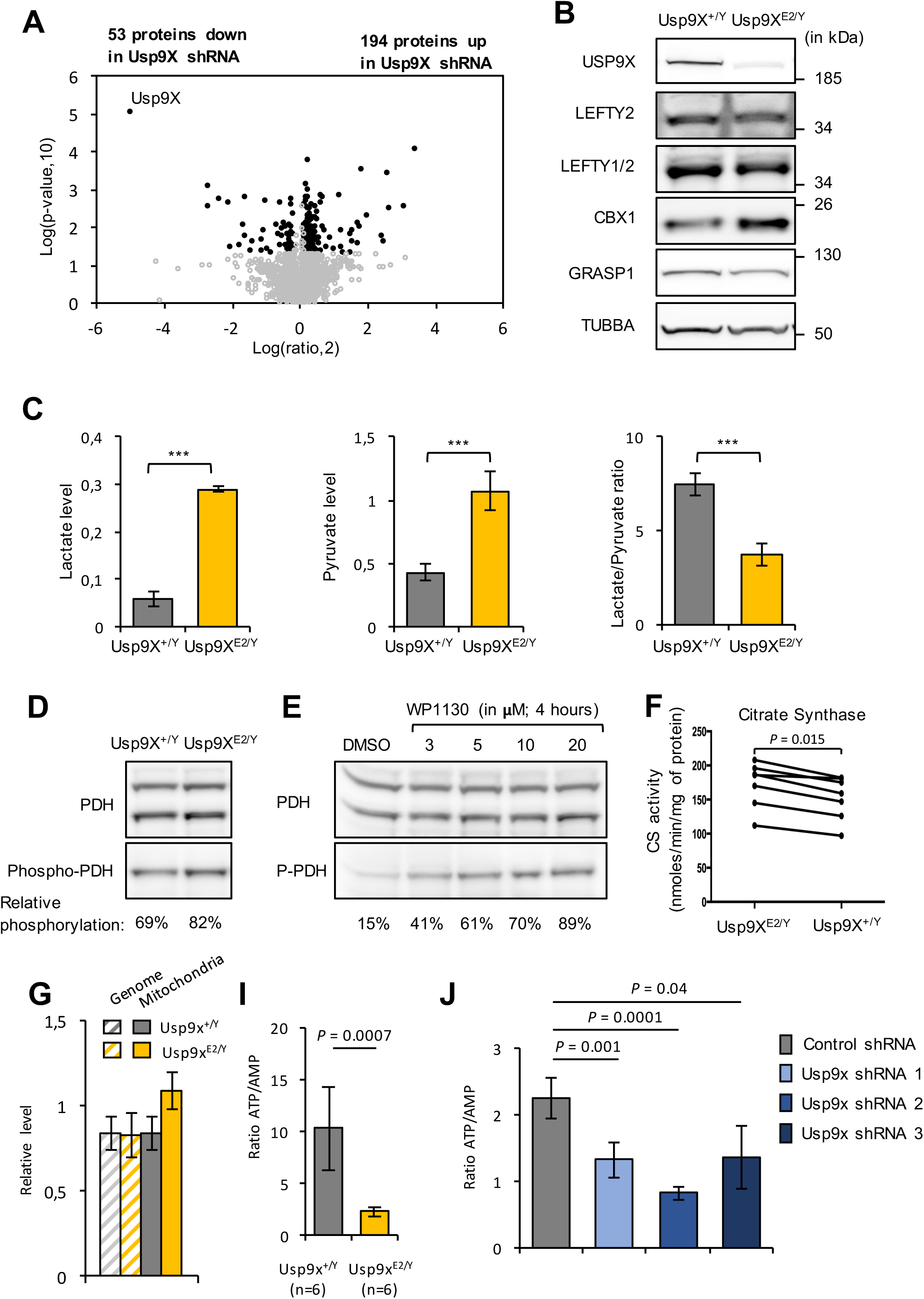
A proteomic analysis unveils a regulation of ATP and glycolysis levels by USP9X. (**A**) Volcano plot showing the differential abundance of proteins in Usp9X^E2/Y^ and control cells. Each point represents the average value of one protein in three replicate experiments. The difference is considered significant for a fold change of ≤-1.1 or ≥1.1 and a *p*-value < 0.05. Proteins meeting these criteria are depicted in black. (**B**) Validation of mass-spectrometry data by western blotting using α-TUBULIN as control. Representative western blots for LEFTY1/2, CBX1, GRASP1 and USP9X (validation of ESRRB is presented in Figure 4G). (**C**) From left to right, lactate levels, pyruvate levels and lactate/pyruvate ratio in the media of Usp9X^E2/Y^ and control ESC grown in LIF + serum. Metabolite levels were measured in 3 independent experiments and data reported as mole per cell. Lactate/pyruvate ratio is presented as mean (± SD) of independent ESC cultures (n = 3). (**D**) Western blot analysis of pyruvate dehydrogenase (PDH) levels and phosphorylation (serine residue 293) in Usp9X^E2/Y^ and control ESC grown in LIF + serum. Relative levels of phospho-PDH (inactive form) / total PDH are indicated. (**E**) Representative western blots of PDH levels and phosphorylation in ESC treated with increasing concentration (3, 5, 10, 20 μM) of WP1130 for 4 hours or vehicle (DMSO). Relative levels of phospho-PDH (inactive form) / total PDH are indicated. (**F**) Measure of citrate synthase activity in Usp9X^E2/Y^ and control ESC (n = 6). *, *P* < 0.05 (paired t-test between groups). (**G**) qPCR quantification of mitochondrial and genomic DNA amounts in Usp9X^E2/Y^ and control ESC. (**H**) Reduction in ATP/AMP ratio in Usp9X^E2/Y^ compared to control ESC. Nucleotides were measured by HPLC. The data presented are mean (± SD) of independent ESC cultures (n = 6). (**I**) Reduction in ATP/AMP ratio in ESC transfected with Usp9X shRNAs compared to control shRNAs (n=3).

We then investigated the involved biological pathways using PantherDB and Ingenuity Pathway Analysis (IPA) for annotation. An equal proportion of misregulated proteins were located in the cytoplasm and in the nucleus (Supplementary Table 3A). We confirmed previous studies showing USP9X involvement in mTORC2/RICTOR and TGFβ pathway regulation in stem cell populations (Supplementary Table 3B) [37,49–52]. IPA software further predicted an inhibition of RICTOR pathway and a down-regulation of differentiation factor GATA6 pathway in Usp9X^KD^ ESC (Supplementary Table 3B). Strikingly, numerous annotations in the PantherDB and Ingenuity Pathway analysis referred to ATP, glycolysis and mitochondria related functions: “mitochondrial dysfunction” (*P* = 1.28e-3), “carbohydrate metabolism” (*P* = 3.86e-2 – 1.1e-4), “oxidative phosphorylation” (*P* = 3.31e-4) (Supplementary Table 3C-D). Considering only up-regulated proteins in the analysis even strengthened the association (Supplementary Table 3E-G). This proteomic analysis indicates a pronounced alteration of ATP pathways and metabolic routes, in Usp9X^E2/Y^ ESC.

We then investigated if these proteomic changes were functionally relevant. Lactate and pyruvate levels were higher in Usp9X^E2/Y^ ESC culture media compared to control ESC, inducing acidification of the culture medium, while the lactate/pyruvate ratio was lower (Figure 2C). Pyruvate dehydrogenase (PDH) phosphorylation was significantly increased in Usp9X^E2/Y^ and upon inhibition of USP9X catalytic activity using WP1130 (or Degrasyn) treatment, indicating its inhibition (Figure 2D-E) [53]. These data indicated that the higher levels of glycolytic enzymes in Usp9X^KD^ ESC are correlated with higher glycolytic activity and lactate production [18,21,23]. The oxidized redox status, shown by the decreased lactate/pyruvate ratio, suggested efficient oxidative phosphorylation pathways. However, the decreased routing of pyruvate towards tricarboxylic acid cycle induced by the inhibition of PDH might, at least partly, underlie that apparent efficacy.

The significant increase in the abundance of proteins involved in oxidation/reduction reactions at the mitochondrial membrane suggested direct mitochondrial modulation (Supplementary Table 1). Furthermore, measurement of citrate synthase activity, a tricarboxylic acids cycle enzyme often used as a proxy for mitochondrial mass, showed significant increase in Usp9X^E2/Y^ ESC as compared to control ESC (Figure 2F). In addition, PCR quantification of the mitochondrial genome also showed a significant increase in Usp9X^E2/Y^ ESC compared to control ESC (Figure 2G). Thus, a mild but consistent increase in the levels of oxidation/reduction proteins abundance is significantly correlated with increased mitochondrial levels in Usp9X^E2/Y^ ESC. In contrast, the amount of proteins involved in mitochondria biogenesis (PGC1, NRF1, TFAM) or in mitochondria fusion-fission events (DRP1, MFN1, MFN2, OPA1) appeared normal (Supplementary Table 1).

Two different approaches evaluated cellular respiration and gave different results. High resolution respirometry analysed the respiration of cells in suspension in their culture medium. Basal, ATP-producing and maximal respiration of the Usp9X^E2/Y^ ESC appeared normal in these conditions. Seahorse technology allowed analysing cells attached. In these conditions, both basal and ATP-producing respiration appeared decreased in two independent experiments (Supplementary Figure 3A). Measurements of two OXPHOS complexes, succinate ubiquinone oxido-reductase (Complex II) and cytochrome c oxidase (Complex IV) did not show significant differences between Usp9X^E2/Y^ and control ESC (Supplementary Figure 2B). Finally, liquid chromatography (HPLC) analysis revealed altered nucleotide balance with severe decrease in ATP, less severe decrease in ADP and increase in AMP levels upon USP9X depletion (Supplementary Figure 3C-E). As a result, the ATP/AMP ratio, a major marker of the bioenergetics status of the cell, was dramatically reduced in Usp9X^KD^ and Usp9X^E2/Y^ ESC as compared to control ESC (Figure 2H-I).

As a whole, our data indicated that depletion of Usp9X favours glycolysis and lactate production in ESC. It also induces modifications of the mitochondrial bioenergetics, including decreased routing of pyruvate towards its oxidation, increased mitochondrial components, and reduced respiration. The ensuing energetic status of the cells is drastically modified, as shown by the nucleotides balance. It is thus likely that USP9X regulates glycolysis and OXPHOS balance during ESC transition through pluripotency states or to actively promote their transitioning.

### USP9X regulates pluripotency transition and its depletion confers a “2i”-like phenotype

Previous proteomic analyses have reported an up-regulation of proteins involved in glycolysis, lipid metabolism, oxidation-reduction and other metabolic routes in naive pluripotency compared to ESC grown in LIF + serum [18, 54]. Our proteomic observations suggested a reinforcement of pluripotency features in Usp9X^KD^ ESC. To better evaluate the role of USP9X on pluripotency status, we then compared the global transcription prolife of Usp9X^KD^ ESC and control ESC in serum + LIF condition by microarray analysis. Using a *p*-value cut-off of 0.05 and a fold change of 1.2, we identified 311 genes differentially expressed between Usp9X^KD^ and control cells, including 170 genes down-regulated and 141 genes up-regulated in Usp9X^KD^ cells (Figure 3A and Supplementary Table 4A). Although the amplitude of gene expression changes was quite modest, it is reminiscent of what has been observed upon depletion of numerous transcription factors or deubiquitinases in murine and human ESC [9,26,42]. Very little correlation also existed between the proteomic and transcriptomic changes in Usp9x^KD^ ESC (Supplementary Figure 4A-B). A functional annotation of USP9X target genes show a significant enrichment for genes involved in “extracellular space”, “cell metabolism”, “cell differentiation” and “neurogenesis” (Supplementary Table 4B-F). This later term is in accordance with a role of Usp9X in neuron development and physiopathology in the adult mice and in humans [31,33,36,37,39,55].

**Figure 3:**
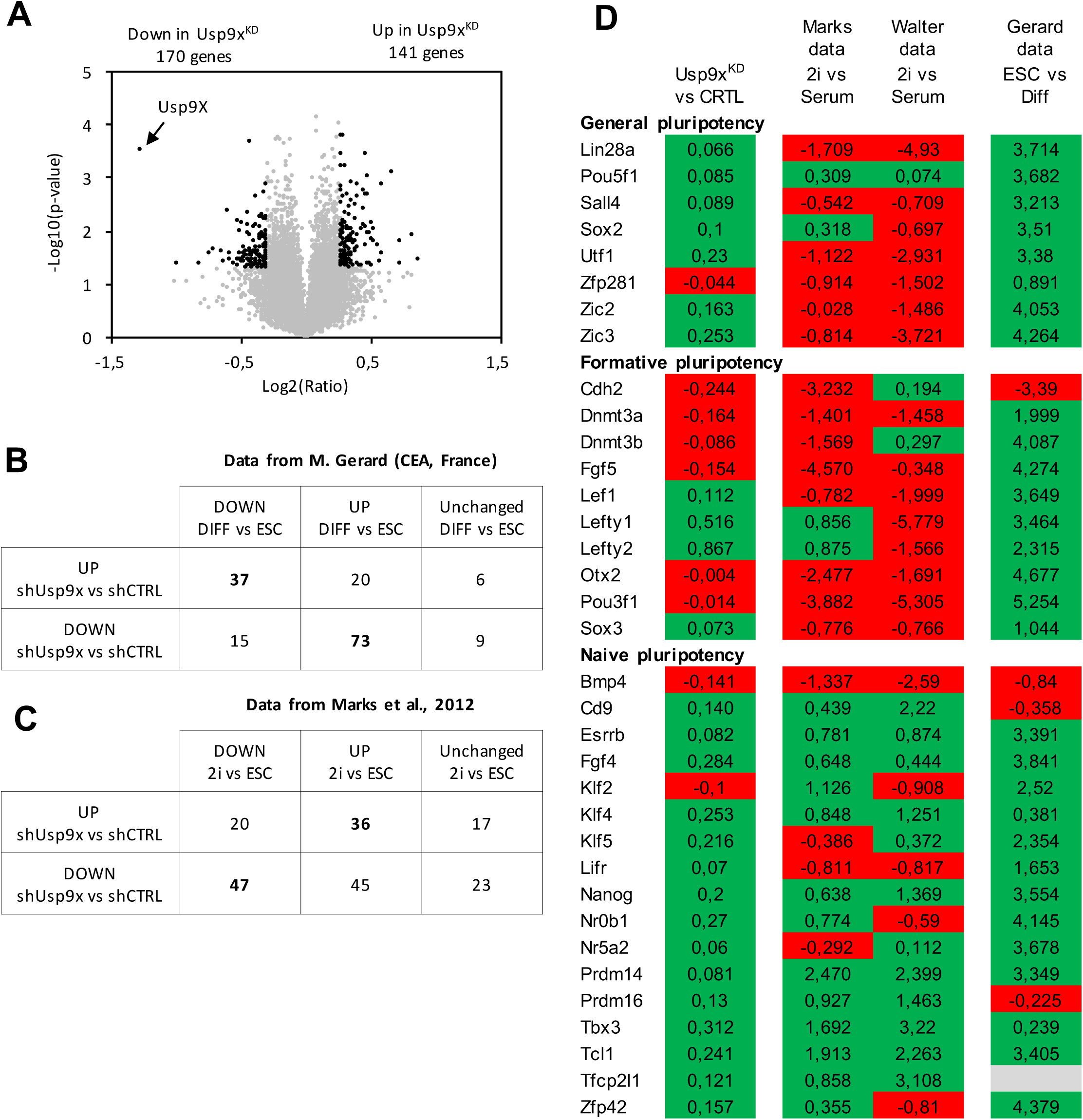
Microarray analysis of USP9X^KD^ ESC. (**A**) Volcano plot displaying differentially expressed genes between ESC transfected with control shRNA and Usp9X-specific shRNAs (Usp9X^KD^ ESC). Each point represents the average value of one transcript in three replicate experiments. The expression difference is considered significant for a fold change of ≤-1.2 or ≥1.2 and a *p*-value < 0.05. Genes meeting these criteria are depicted in black. (**B**) Intersection of differentially expressed genes identified from Usp9X^KD^ ESC (vs control ESC) and a RNA-sequencing dataset of differentially expressed genes during ESC differentiation (data kindly shared by M. Gerard, CEA, France). (**C**) Intersection of differentially expressed genes identified from Usp9X^KD^ ESC (vs control ESC) and array datasets of differentially expressed genes between ESC cultured in LIF + serum or in “2i” (data from Marks et al., 2012). (**D**) Representation of core, formative and naive pluripotency transcription factors expression in Usp9X^KD^ vs control ESC, in 2i vs LIF + serum and in ESC vs Differentiated cells. Positive fold change is represented in green; negative fold change in red. Fold-change values are indicated.

In order to understand how USP9X may act upon target genes, we searched for transcription factor motifs enrichment using the oPOSSUM web-based platform. We revealed that genes deregulated in Usp9X^KD^ cells were enriched in their promoter for DNA motifs of core pluripotency transcription factors, including KLF4, ZFP281 and MYC (Supplementary Table 4G-H) [56–58]. This suggests that USP9X transcriptional targets are also regulated by pluripotency factors. Interestingly, when the Usp9X transcriptional signature was compared to other genetic and chemically perturbed background in the MSigDB collection or to datasets in the StemChecker repository we retrieved further evidence that USP9X regulates expression of genes involved in cell differentiation that are regulated by pluripotency transcription factors (Supplementary Table 4E and 4I).

To determine the transcriptional pluripotency status of Usp9X^KD^ ESC, we investigated if the changes in gene expression observed in Usp9X^KD^ ESC might mirror the kinetics of gene expression of ESC adapted to different pluripotency status or progressing in differentiation. We first used a dataset of gene expression comparing ESC with differentiated cells. We observed that 75.2% of the genes down-regulated in Usp9X^KD^ ESC are expressed at higher levels in differentiated cells compared to ESC grown in LIF + serum (Figure 3B). Conversely, 58.7% of the genes up-regulated in Usp9X^KD^ are expressed at lower levels in differentiated cells compared to ESC grown in LIF + serum (Figure 3B). Similar results were obtained using two other gene expression datasets comparing ESC to differentiated cells (NCBI GEO: GSE96809 and GSE124241) (Supplementary Figure S5A) [59, 60]. These data indicate that Usp9X^KD^ ESC exhibit stronger repression of genes upregulated during differentiation and higher expression of genes downregulated during differentiation, as observed in the naive or “2i” condition.

We thus curated the literature data to identify genes that are over and under expressed in “2i” culture compared to LIF + serum. Using datasets by Marks et al. we observed that 40% of the genes down-regulated in Usp9X^KD^ ESC are expressed at lower levels in “2i” ESC compared to ESC grown in LIF + serum (Figure 3C) [61]. Conversely, 49% of the genes up-regulated in Usp9X^KD^ are expressed at higher levels in “2i” ESC compared to ESC grown in LIF + serum (Figure 3C). A similar trend was observed using datasets from NCBI GEO GSE71591 (Supplementary Figure S5B) [62]. These observations suggested that depletion of USP9X causes transcriptional changes similar to ESC transitioning from LIF + Serum to “2i”.

We thus directly investigated the levels of expression of factors classified as regulators of naive, formative and general pluripotency [14]. We observed that Usp9X^KD^ ESC exhibit an up-regulation (although not statistically significant) of most factors associated with naive pluripotency and general pluripotency (Figure 3D). On the contrary, factors associated with formative (or primed) pluripotency tend to be less expressed. Overall these results, as well as proteomic data, indicated that depletion of Usp9X alters the pluripotency status of ESC by promoting transcriptional and metabolomics features associated with naive pluripotency and that Usp9X regulates the transition of ESC through pluripotency stages.

### USP9X interacts with proteins essential for pluripotency and early embryogenesis

In an attempt to characterize the function of USP9X in ESC we used a single label-free proteomic approach to generate an USP9X interaction profile in ESC (Figure 4A). We successfully immunoprecipitated significant levels of endogenous USP9X from total cellular extracts prepared from ESC maintained in LIF + serum and identified 460 proteins significantly enriched in USP9X IPs compared to control IgG (Figure 4B and Supplementary Table 5A). Most of the putative interactors were not previously reported in similar analyses in human cancer cell models suggesting that they mostly represent ESC-specific interactions. Importantly, we detected in USP9X precipitates, pluripotency transcription factors SALL4 and OCT4 as well as Gag proteins PEG10 and PEG3, as previously reported in ESC (Supplementary Table 5A-B) [63, 64].

**Figure 4:**
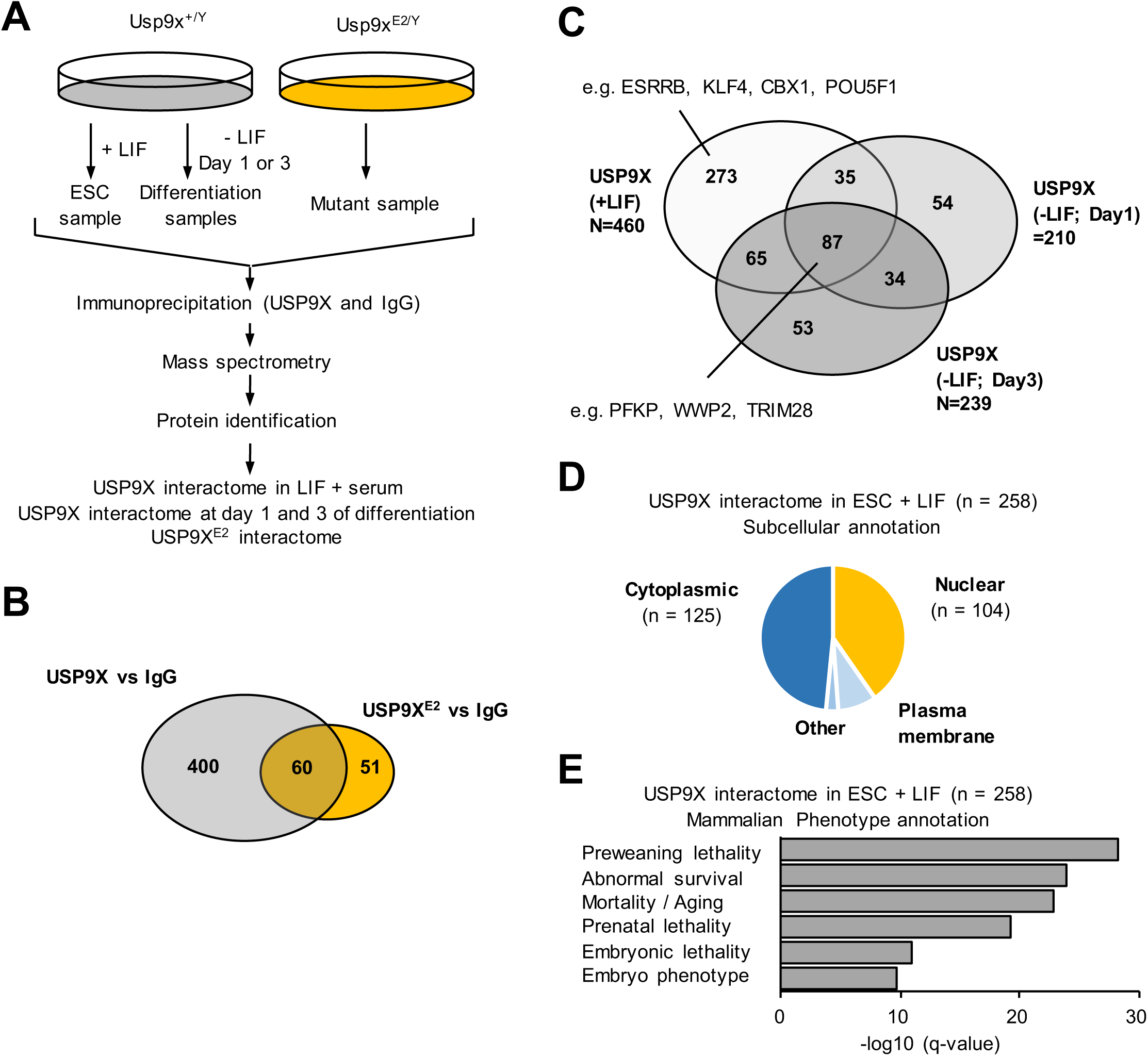
Co-immunoprecipitations reveals distinct interaction partners of USP9X in LIF + serum medium, during differentiation and in the USP9X^E2/Y^ background. (**A**) Schematic diagram of the quantitative MS-based workflow for the identification of potential USP9X binding partners in LIF + serum medium and upon differentiation (at day 1 and day 3), as well as potential binding partners of USP9X^E2^. The experimental workflow included protein extraction, immunoprecipitation with a specific anti-USP9X antibody or IgG as control, tryptic digestion, mass-spectrometry and peptide identification. (**B**) Venn diagram analysis of common and specific partners of USP9X and USP9X^E2^ in LIF + serum. (**C**) Venn diagram analysis of common and specific partners of USP9X in LIF + serum and in day 1 and 3 of differentiation. (**D**) Chart of the subcellular localization of the USP9X-interacting proteins detected solely in LIF + serum. (**E**) Classification of USP9X-interacting proteins in LIF + serum according to phenotypic annotations.

As a control we also performed an immunoprecipitation with USP9X antibody in USP9X^E2/Y^ ESC. We identified 110 proteins with 60 proteins common to the USP9X interactome and 51 new proteins (Figure 4B and Supplementary Table 5C). We thus identified the network of USP9X binding partners at endogenous expression levels in ESC and most of these interaction partners were not detectable in the Usp9X^E2/Y^ background.

We then generated the USP9X interaction profiles at days 1 and 3 after LIF removal and induction of cell differentiation. We identified 210 and 239 proteins co-purifying with USP9X at day 1 and day 3 of differentiation respectively, including PEG3 and PEG10 (Figure 4C and Supplementary Table 5D-E). An intersection of these different lists (including USP9X^E2^ partners), identified a list of 258 proteins constituting the USP9X-specific interactome in ESC (Supplementary Table 5F-G).

An annotation of these 258 potential binding partners indicated a roughly equal number of proteins located in the cytoplasm (n = 125) and in the nucleus (n = 104) (Figure 4D). A functional annotation revealed an enrichment for molecular functions such as protein-, nucleic acid- and ATP-binding and for biological functions such as metabolic processes, translation and biosynthesis (Supplementary Table 6A-E). Importantly, it also revealed an enrichment for genes essential for early embryonic development (Figure 4E and Supplementary Table 6A-B). An annotation with the REACTOME database showed similar enriched functions whether we considered the 460 potential interactors in LIF + serum or solely the 258 proteins constituting the USP9X-specific interactome in ESC (Supplementary Table 6F). On the contrary, potential USP9X interactors detected in LIF + serum as well as during differentiation (n = 87) are mostly enriched for generic cellular functions such as cell cycle, DNA synthesis and metabolism (Supplementary Table 6G).

In sum, the characterisation of USP9X interactome shows that several of USP9X ESC-specific potential binding partners are involved in cell metabolism and gene expression regulation, as anticipated from functional assays. Furthermore, some of them were previously reported as regulators of (ESC and human cancer) stem cell identity, including chromatin factor CBX1 and pluripotency factors OCT4, KLF4 and ESRRB [5,6,65]. However, USP9X has not been reported to regulate their function in human cancer cells or murine ESC.

### USP9X regulates ESRRB abundance and transcriptional activity

We then reasoned that USP9X might regulate the function of naive pluripotency transcription network. This hypothesis was further supported by the observation that transcription factor ESRRB regulates OXPHOS and glycolysis balance during cell reprogramming [66, 67]. We thus investigated whether USP9X interacted with ESRRB in ESC, as suggested from our proteomic analysis (Supplementary Table 5). We performed co-immunoprecipitation assays on total ESC cellular extract. We immunoprecipitated significant amount of ESRRB and detected by immunoblotting USP9X in ESRRB co-immunoprecipitates (Figure 4A). These data validated that USP9X binds ESRRB in ESC.

We then investigated whether USP9X regulates ESRRB abundance. We performed ESRRB-staining coupled with FACS analysis. We observed that ESRRB levels were higher in Usp9X^E2/Y^ ESC compared to control cells when cells are cultured in LIF + serum (Figure 2B). On the contrary, when ESC were adapted to “2i”, ESRRB levels were high in Usp9X^E2/Y^ ESC and control cells, with no difference between levels detected in both cell types (Figure 2B). Surprisingly, USP9X levels were lower in 2i compared to LIF + serum in similar samples (Supplementary Figure 4D). Finally, we confirmed ESRRB regulation using alternative approach. We observed higher levels of ESRRB in Usp9X^E2/Y^ ESC compared to control cells in LIF + serum by western blot (Figure 2C). We also observed higher levels of ESRRB in ESC transfected with Usp9X shRNAs compared to control ESC (Supplementary Figure 4E). These data indicate that depletion of USP9X up-regulates ESRRB levels in ESC.

To understand the underlying mechanisms of USP9X depletion on ESRRB function, we investigated whether ESRRB was ubiquitinated. We immunoprecipitated ESRRB from ESC treated with MG132 (10 mM for 4 hours) or not and analysed the samples by mass-spectrometry. We detected significant amount of ESRRB in both IPs (as well as USP9X) and obtained similar coverage (58%) of ESRRB in both samples (Supplementary Figure 4A). We then searched for ESRRB peptides containing an ubiquitinated amino-acid. We identified a peptide (TIQGNIEYNCPATNECEITKR) spanning amino-acids 130 to 150 of ESRRB with an ubiquitin modification (Supplementary Figure 4B). To confirm ubiquitination of ESRRB, we probed ESRBB immunoprecipitates using an ubiquitin-specific antibody. We observed an ubiquitin signal at a molecular mass consistent of ESRRB being mono-ubiquitinated (Figure 5D). These data confirmed that ESRBB is a ubiquitinated protein in ESC cells and identified a ubiquitination site not previously reported [68].

**Figure 5:**
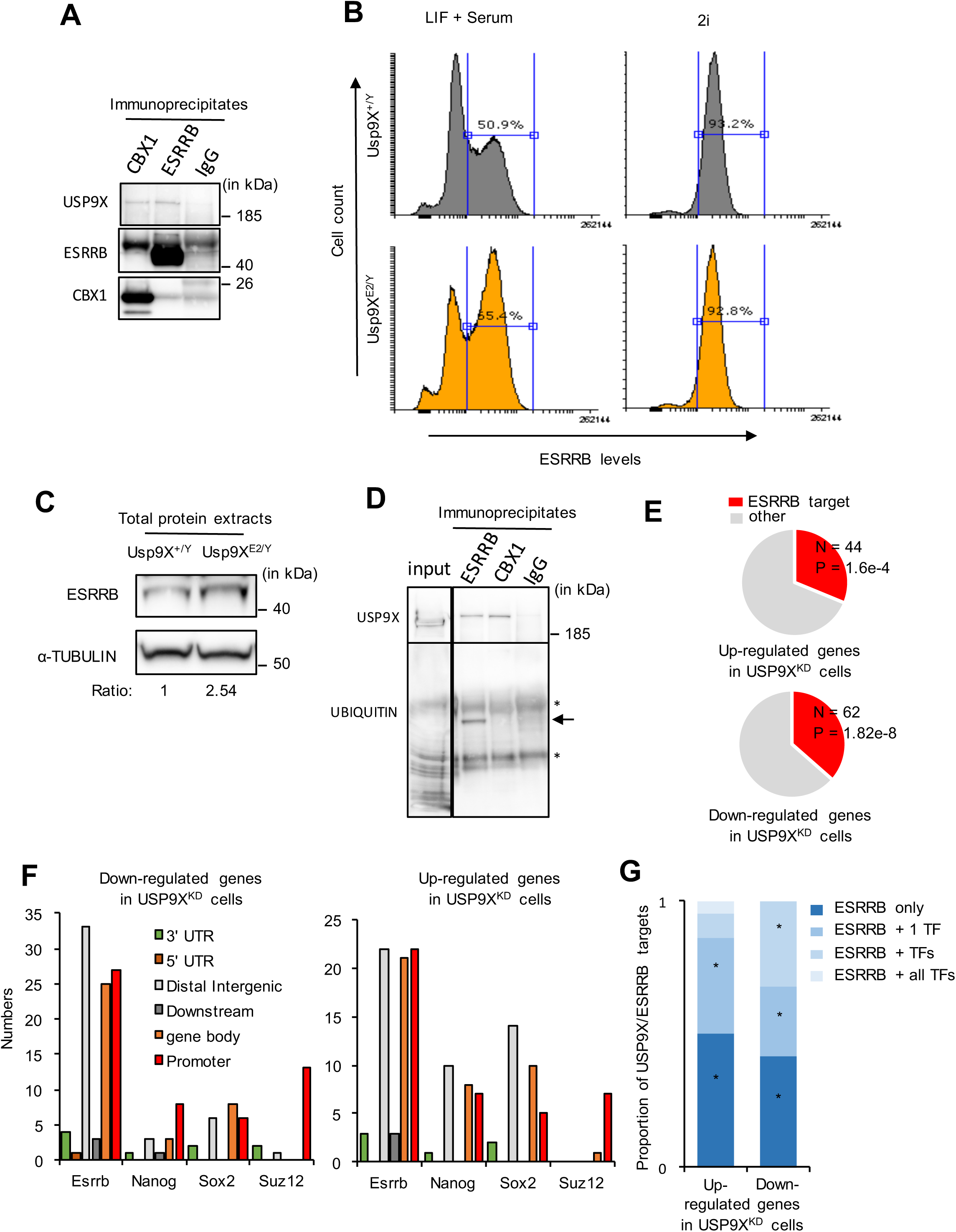
USP9X interacts with ESRRB and regulates its abundance and function. (**A**) USP9X co-immunoprecipitates with ESRRB and CBX1 but not irrelevant IgG in total cellular extracts prepared from ESC grown in LIF + serum. (**B**) Flow cytometry analysis of ESRRB protein levels in Usp9X^E2/Y^ and Usp9X^+/y^ ESC cultured in LIF + serum (on the left) or “2i” (on the right). Hundred thousand cells were analysed per condition. (**C**) Western blot analysis of ESRRB and α-TUBULIN abundance in Usp9X^E2/Y^ and Usp9X^+/y^ ESC cultured in LIF + serum. The relative ESRRB/α-TUBULIN ratio is indicated under the blots. (**D**) Cells grown in LIF + serum were treated with MG132 (10mM for 4 hours) and then lysed for immunoprecipitation using an anti-ESRRB, an anti-CBX1 and control irrelevant IgG. The immunoprecipitates were further probed for the levels of ubiquitination of ESRRB and CBX1 using anti-ubiquitin antibody. The expected size of ESRRB is indicated by an arrow. *, immunoglobin heavy and light chains. (**E**) Overlap between genes up and downregulated upon Usp9X depletion and genes regulated upon Esrrb depletion (data from Nishiyama et al. 2013). (**F**) Annotation of Usp9X transcriptional targets according to the presence of ESRRB, NANOG, SOX2 or SUZ12 binding sites in their vicinity. Binding sites were classified according to their genomic annotation: 3’-UTR, 5’-UTR, Distal Intergenic, Downstream (immediate downstream of a gene, within 3 kilo-bases), Gene body and Promoter (or transcription start site) of Usp9X regulated genes. (**G**) Proportion of Usp9X/Esrrb regulated genes (up or down) also transcriptionally regulated by an additional or combination of pluripotency factors (i.e. were considered for this analysis: Nanog, Sox2, Oct4, Nr0b1/Dax1 and Sall4 datasets from Nishiyama et al. 2013) (*, *p* < 0.01, Hypergeometric test).

We then pre-treated the cells with proteasome inhibitor MG132 (10 mM for 4 hours) prior to ESRRB immunoprecipitation. We observed that MG132 treatment does not abrogate the interaction between USP9X and ESRRB by western blot and mass-spectrometry (Supplementary Figure 4C). It also does not cause the appearance of poly-ubiquitination signals in ESRRB IPs (Supplementary Figure 4C). We then investigated the role of USP9X deubiquitinase activity on ESRRB ubiquitination by utilizing WP1130, the USP9X-catalytic inhibitor [53]. We co-treated ESC with MG132 and WP1130 prior to ESRRB immunoprecipitation. We did not observe significant changes in ubiquitination levels of ESRRB upon addition of WP1130 (Supplementary Figure 4C). These data indicate that depletion of Usp9X does not change ESRRB protein ubiquitination levels in our assay, but does not rule out that USP9X may regulate ESRRB transcriptional activity.

Using previously reported transcriptomic analysis (NCBI GEO GSE26520) we observed that a significant fraction of USP9X target genes are transcriptional targets of ESRRB (Figure 5E). We thus intersected gene expression data from Usp9X^KD^ ESC with available chromatin immuno-precipitation data for ESRRB and other factors important for ESC pluripotency and differentiation (NCBI GEO GSE11431) [69]. We observed that a large proportion of USP9X target genes, either up or downregulated, were bound at their promoter (or gene body) by ESRRB and to a lesser extent by the pluripotency factors NANOG and SOX2, and by the Polycomb protein SUZ12 (Figure 5F). These data suggested that ESRRB and USP9X directly co-regulates a subset of genes essential to define ESC pluripotency (Figure 5F). Importantly, this transcriptional program appears independent from the activity of other pluripotency transcription factors for the most part (Figure 5G). It is thus likely that USP9X binding to ESRRB destabilizes the protein or its interaction with other pluripotency factors facilitating the transition through ESC states, and changes in cell metabolism.

## Discussion

### USP9X regulates the activity of the pluripotency network

It has been repeatedly observed that USP9X is highly expressed in stem cells (from different tissues) and that its expression is lower in differentiated cells. It is thus often extrapolated from these mRNA expression studies that USP9X is important to define pluripotency [4, 41]. Our work clearly demonstrates that depletion of USP9X by shRNA or removal of exon 2 is actually not sufficient to trigger spontaneous ESC differentiation and the loss of pluripotency features. Depletion of USP9X actually causes an imbalance in pluripotency transcription factor expression and abundance, such as LEFTY2 (a regulator of TGFβ signalling) and (naive) pluripotency factors ESRRB and KLF4. This imbalance in pluripotency factors has functional consequences. First, depletion of USP9X facilitates cell differentiation in embryonic bodies and extra-embryonic stem cells (XEN) upon permissive culture conditions. Second, depletion of USP9X modifies cell metabolism and the activity of signalling pathways associated with naive (or 2i) ESC.

Intriguingly, different studies, with variable conclusions, have reported a function of USP9X in TGFβ pathway during early embryogenesis, in adult neurons, in cancer cells and in Drosophila tissues [36,37,49,50,70]. Our data suggest an additional function in ESC, with a new layer of regulation involving the regulation of LEFTY2 abundance, a secreted ligand that may act through cell autonomous and non-autonomous functions to regulate ESC fate. We also confirmed previous studies indicating a function of USP9X in mTOR signalling in stem/progenitor cells and its interaction with PEG proteins [51,52,63].

Our multi-omic analyses demonstrate that depletion of Usp9X actually reinforces pluripotency, with higher levels of “naive” transcription factors ESRRB, higher differentiation potential, higher glycolytic activity and higher lactate production. These observations are not consistent with a previous study showing that depletion of Usp9X by siRNAs causes reduction of an endogenous GFP-tagged NANOG protein and thus loss of pluripotency [26]. This discrepancy may be the consequence of technical differences between the studies. Among many things, the nature of ESC studied, the levels of USP9X depletion obtained, off-target effects and the culture condition (feeders in our study vs gelatin in Buckley et al). may alter the response of ESC to Usp9X depletion [20, 71]. We noted variable effects of Usp9X depletion on mRNA and protein expression suggesting it may be challenging to solely rely on transcriptomic analysis and single reporter gene constructs to biologically characterized USP9X function in ESC. In line with this observation, it was recently demonstrated that LEFTY2 mRNA expression increases upon withdrawal of LIF while the protein level is dramatically reduced, as we observed in Usp9X^E2/Y^ ESC [15, 72].

Furthermore, many genes regulated by USP9X at mRNA levels are actually constituents of the extracellular compartment (72 out of 311 genes, including LEFTY2), and thus probably not properly detected in our proteomic analysis. These constituents of the niche might alter ESC fate. To date, few studies are investigating the contribution of the niche on ESC biology, needless to say at the transition between pluripotency states. Our proteomic and transcriptomic analysis provide an extensive resource to investigate additional functions of USP9X in ESC.

### USP9X regulates the transition of ESC through pluripotency states

Both proteomic and functional assays unveiled that USP9X regulates ESC cell metabolism. We observed increased levels of lactate, pyruvate and decrease levels of ATP. These molecules are important cellular and extra-cellular messengers in ESC [73]. For instance, higher levels of lactate production and release may cause alteration in histone acetylation and lactilation levels in ESC with in turn may interfere with gene expression and chromatin organization [74].

These metabolic changes, as well as the transcriptomic changes, observed in Usp9X^E2/Y^ and Usp9X^KD^ ESC are consistent with ESC exhibiting a more naive ESC status. Intriguingly, EpiSC, that are more developmentally advance, also show an increase in glycolytic activity and lactate production compared to ESC grown in LIF + serum [75]. The coordination between the kinetic of expression of pluripotency factors and the balance between OXPHOS and glycolytic metabolism might controls ESC transition from naive to EpiSC. USP9X^E2/Y^ ESC may thus represent a transition step characterized by a strong repression of differentiation genes, higher levels of naive pluripotency factors and higher glycolysis. Our data are thus generally consistent with a role of USP9X in the transition between states of pluripotency by regulating a subset of pluripotency target genes and cell metabolism.

### USP9X regulates ESRRB function

Co-immunoprecipitation assays followed by proteomic identified numerous proteins interacting with USP9X in ESC, and most of these interaction are disrupted by the deletion of exon 2 of Usp9X or when cells enter differentiation upon LIF removal. A recent study reported USP9X potential binding partners in ESC using a flag-tag inserted in frame in Usp9X exon 2 [63]. The study reported far less USP9X potential binding partners compared to our work, most of which are detected in the Usp9X^E2/Y^ background. It is thus tempting to speculate that insertion of the Flag-tag in Usp9X exon 2 creates a loss-of-function mutant, and thus explains the relative low number of potential USP9X partners detected in this study. It also suggest that further work will be needed to investigate whether the extreme N-terminus of USP9X regulates its function in ESC.

While many naive transcription factors interact with USP9X in ESC, our transcriptomic analysis revealed a strong overlap with ESRRB transcriptional targets. By western blot analysis we revealed that USP9X binds to ESRRB in ESC and that depletion of USP9X causes an up-regulation of ESRRB protein with no significant changes in Esrrb mRNA levels. Intriguingly, ESRRB is a substrate of ubiquitin, glutamylation and phosphorylation [68]. Our proteomic analysis could not detect peptides overlapping the glutamylation sites (Serine 25) nor the previously described ubiquitination site [68]. We however notice a ubiquitination site on a peptide spanning amino-acid 130 to 150 of ESRRB and including a lysine or cysteine as potential modified sites. Importantly, ESRRB ubiquitination appears to be mono-ubiquitination as revealed by western blot analysis. So far, our data suggest that ubiquitination of ESRRB is not affected by depletion of Usp9X or treatments with WP1130, a USP9X DUB inhibitor. It is thus possible that ESRRB ubiquitination favour its interaction with USP9X, that in turn increases its stability through protein/protein interactions and its function at transcriptional targets.

Intriguingly, Esrrb knock-out ESC did not show dramatic morphological and transcriptional changes [9, 66]. Nonetheless, ESRRB controls cell metabolism and energy balance in pluripotent cells and during cell reprogramming [20,66,67,76]. It is thus possible that cell metabolism changes observed upon Usp9X depletion are a consequence of ESRRB transcriptional activity. It has also been shown that ESRRB acts as a bookmarking factor important to re-established the naive pluripotency transcriptional program when ESC enter the next the next G1 phase of the cell cycle [76].

Alternatively, many metabolic enzymes and regulators of cell metabolism were detected in Usp9X immunoprecipitates, including glycolytic enzymes. We can thus also propose that USP9X regulation of metabolic routes may alter the energy balance in ESC, fine tunes signalling pathways activities and promotes changes in the pluripotency network composition and activity. Indeed, OXPHOS and glycolytic metabolisms promptly evolve as ESC transition through pluripotency states.

Finally, analysis of available ChIP-sequencing datasets indicates that the promoter of Usp9X is occupied by OCT4, NANOG, SOX2, KLF4, ESRRB and NR5A2, as previously reported in the BindDB database [77]. Consistent with Usp9X being a transcriptional target of the pluripotency network, we observed that USP9X is down regulated with a similar kinetics as pluripotency factors upon differentiation in embryonic bodies. It is thus likely that USP9X is a direct transcriptional target of the pluripotency network but also a regulator of the activity of this network at specific genes.

While mRNA studies suggested a function of Usp9X in governing pluripotency, our data suggest a more complex function. Contrary to expectation, depletion of Usp9X does not cause spontaneous differentiation but rather reinforce pluripotency features. Through the regulation of cell metabolism, transcription factor abundance and other ESC features, Usp9X regulates the transition of ESC through pluripotency states, and its depletion enhances the developmental potential of ESC. These findings may have important implications in the design of the cocktails utilized for somatic cell reprogramming.

## Materials and methods

### Mouse embryonic stem cells

Mouse male embryonic stem cells (ESC) V6.5 were obtained from Pr Kian Peng Koh (KU Leuven, Belgium). Usp9X^E2/Y^ and control Usp9X^+/y^ male ESC were previously described and derived from C57BL/6N mice [45].

ESC were grown at 37°C with 5% CO2 in a humidified incubator. ESC were cultured on feeders using the Dulbecco’s Modified Eagle’s Medium (DMEM) (High Glucose and Pyruvate) supplemented with 15% fetal bovine serum, 1% L-Glutamine (Gibco, 25030-081), 1% Penicillin/Streptomycin (Life Technologies, 15140163), 100 µM β-mercaptoethanol (Sigma-Aldrich, M7522) and 500 U/ml leukemia inhibitory factor (Chemico, ESG1107). Mouse embryonic fibroblast were obtained at P0 and cultivated in DMEM (High Glucose and Pyruvate) plus 10% serum until P4 then treated with 10 µg/mL mitomycin C (Sigma-Aldrich; M428).

Cells were also grown in feeder free conditions using « 2i » media consisting of 50% DMEM-F12 (Thermofisher, 10565018), 50% Neurobasal (Thermofisher, 21103049), supplemented with 0.5 mM L-glutamine (Gibco, 25030-081), 1% Penicillin/Streptomycin (Life Technologies, 15140163), 1X nonessential amino acids (Gibco, 11140-050), 0.1mM β-mercaptoethanol (Sigma, M7522), 1000 U/ml LIF (Chemico, ESG1107), B27 supplement (Thermofisher, 17504044), N2 supplement (Thermofisher, 17502048), 1µM PD0325901 (Cell Guidance Systems, SM26-2) and 3µM CHIR99021 (Cell Guidance Systems, SM13-1) [78].

Differentiation of ESC into embryonic bodies was performed as previously described [46]. 4 x 10^6^ cells were cultivated in non-adherent petri dishes (Corning, 351029) and retinoic acid (BioTechne, 0695) added at day 4 and 6 of differentiation at 5 µM final concentration [46]. Differentiation of ESC into extra-embryonic stem cell (XEN) was performed as previously described [47].

WP1130 (or Degrasyn) (Calbiochem, 681685) and InSolution MG132 (Calbiochem, 474791) were purchased from Sigma-Aldrich and utilized at 5 µM and 10 mM final concentration for 4 hours respectively.

### shRNA plasmids and electroporation

Short-hairpin (sh) RNA were constructed into the pHYPER vector as previously described [79]. shRNA sequences targeting Usp9X are described in Supplementary Table S7. Control shRNA (or linker shRNA) correspond to the empty pHYPER plasmid. An additional control plasmid targeting the Icos gene, silenced in ESC, was also contructed and shRNA sequences described in Supplementary Table 7.

Plasmids were electroporated in ESC using the Lonza Nucleofector^TM^ 2B Device using program A-23 (Lonza, AAB-1001). 10 µg of plasmids were mixed with 5 millions ESC in buffer “Mouse ES cell nucleofector kit (Lonza, VPH-1001) per electroporation. 16 hours post-electroporation, puromycin (2 µg/mL) was added to the culture for 48 hours.

### Proliferation assay

Cells were seeded in 6-well plates plated with 0,1% gelatin and MEFs when indicated (Sigma-Aldrich, G18990). The number of cells from duplicate plates were counted manually with a hemocytometer every day. Cell numbers were then plotted relative to the number of cells at day 1.

### Alcaline Phosphatase staining

Alcaline phosphatase activity was determined using a detection kit according to manufacturer recommendations (Sigma-Aldrich, SCR004).

### Western blotting procedure

Standard western blotting was conducted as previously described [80]. Cells were lysed and sonicated in Pierce RIPA SDS buffer (ThermoFisher Scientific, 89901) with protease inhibitor (phenylmethylsulfonyl fluoride and cOmplete^TM^ protease inhibitor cocktail tablets) for 1 hour on ice with 1 µL of benzonase (Sigma-Aldrich, E1014) per ten million cells. Equal amounts of protein were subjected to SDS-PAGE using pre-cast 4-12% BOLT Bis-Tris Plus gels (ThermoFisher Scientific) and SDS Running MOPS buffer (ThermoFisher Scientific, B0001). After protein separation by electrophoresis, samples were transferred to an Immobilon-P PVDF membrane (Sigma-Aldrich, IPVH00005) followed by immunoblotting with the antibodies. Detection was performed using the Super Signal West Dura chemiluminescence horseradish peroxidase substrate (ThermoFisher Scientific, 34076). A list of primary and secondary antibodies is available in Supplementary Table 7.

### RNA extraction, reverse transcription and qRT-PCR analysis

RNA were solubilized and extracted using TRIzol (Ambion, 15596026) as described by the manufacturer. DNA contaminants were removed using the DNA-free^TM^ kit DNAse treatment and removal following manufacturer recommendation (Ambion, AM1906). RNA quality and quantity was then evaluated using a Nanodrop^TM^ 2000 spectrophotometer (Ozyme) or the Agilent 2100 Bioanalyzer system (Agilent).

Reverse transcription was performed using SuperScript^TM^ III or IV reverse transcriptases following manufacturer’s guidelines (references 18080044 and 18090050); ThermoFisher Scientific) and random hexamers (New England Biolabs, S1230S). qPCRs were performed on a Viia 7 real-time PCR system (ThermoFischer scientific) available on the “Epigenomics Core Facility” (UMR7216, Paris) with SYBR Green mix. Relative quantification using a standard curve method was performed for each gene and Gapdh or Ppia were choosen as reference genes. A list of primers is available in Supplementary Table 7.

### Immunofluorescence microscopy

For immunofluorescence staining, cells were prepared as previously described [81] with minor modifications. Briefly, ESC were cultured on 0.1% gelatinized tissue culture plates for 24 hours in order to get rid of the feeders, trypsinized, washed in PBS, adhered to poly-L lysine-coated coverslips and centrifuged for 1 min at 500 RPM at room temperature. Cells were then washed in PBS and subsequently fixed with 4% PFA diluted in PBS for 10 minutes on ice and treated with ice cold 0.5% Triton X-100/PBS for 5 minutes. Cells were rinsed twice with ice cold PBS and blocked in 1% BSA for 30 minutes at RT. Cells were incubated with primary anti-USP9X antibody (Bethyl Laboratories, A301-351A, 1:300 dilution) in 1%BSA for 1 hour at RT. After 3 rounds of PBS washes, the primary antibody was recognized by secondary antibody (A32732; Goat anti-Rabbit IgG (H+L) Highly cross adsorbed Alexa Fluor Plus 555, Thermo Fisher Scientific, dilution 1:500) for 45 minutes in the dark at RT. Cells were finally rinsed twice with PBS for 5 minutes at RT and mounted on ProLong Gold anti-fade reagent supplemented with DAPI (ThermoFisher Scientific, P36931). Images were acquired at the Olympus BX63F equipped with a Hamamatsu ORCA-Flash4.0 LT C11440-42U (CMOS) on a 63x objective using MetaMorph acquisition software. Images were post-processed using Fiji Is Just ImageJ (FIJI) [82].

### Flow cytometry

Flow cytometric analyses were performed on single cell suspension (1 x 10^6^ cells) and dead cells were excluded with Live/Dead staining (eBioscience, 65-0865-14). Intracellular detection of ESRRB and USP9X protein were performed on fixed and permeabilized cells using Intracellular Fixation and Permeabilization Buffer kit (eBioscience), following manufacturer’s instructions with minor modifications. Briefly cells were resuspended in 1 mL of Fixation/Permeabilization working buffer (eBioscience), instantly vortexed and incubated for 60 minutes at 4°C. Cells were then centrifuged at 400g for 5 minutes at room temperature, resuspended in 2 mL of Permeabilization Buffer (eBioscience) and centrifuged again at 400g for 5 minutes at room temperature. After discarding the supernatant cells were resuspended in 100 µL of Permeabilization Buffer containing the antibody of interest (anti-USP9X, 1:300 dilution or anti-ESRRB, 1:500 dilution) and incubated for 1 hour at 4°C. Cells were then rinsed with 2 mL of Permeabilization Buffer and centrifuged at 400g for 5 minutes at room temperature. Supernatant was again discarded and cells were resuspended in 100 µL of Permeabilization Buffer containing the secondary antibody (Anti-rabbit IgG H&L Cy-5, ab97077) and incubated overnight at 4°C. On the next day, cells were twice rinsed with 2 mL of Permeabilization Buffer and centrifuged at 400g for 5 minutes at room temperature. After discarding the supernatant cells were finally resuspended in Flow Cytometry Staining Buffer and analysed on a LSRII flow cytometer (BD Biosciences), equipped with FACS Diva software (BD Biosciences).

### Cell cycle analysis

Edu or 5-ethynyl-2’-deoxyuridine was added to the cells at 10 μM for 20 minutes prior to harvesting. Cells were then fixed, permeabilised, blocked and stained using the Click-IT^TM^ EdU Alexa Fluor^TM^ 488 flow cytometry assay kit (ThermoFisher Scientific, C10425) according to manufacturer recommendation. Samples were further stained with 7-AAD or 7-aminoactinomycine (AAT Bioquest, 17501). Data were acquired on a BD Accuri^TM^ C6 flow cytometer (BD Biosciences) and analyzed using the manufacturer software based on Edu and 7-AAD profiles. Ten thousand cells were analysed per experimental condition. Comparison between groups was performed using Z-score for two population proportion. *P* < 0.05 were considered statistically significant.

### Replication timing analysis

ESC transfected with Usp9X and control shRNA were pulse-labelled with bromodeoxyuridine. Replication timing analysis was then performed and analysed as previously described [83].

### Teratoma analysis

ESC were harvested at 80% confluence by incubation in stem cell trypsin (0.25% trypsin, 0.02% EDTA, 0.1% glucose, 0.3% Tris-HCl in PBS) for several minutes at 37 °C until dissociation of stem cell colonies occurred. The colonies were harvested before MEFs detached. The ESC were washed subsequently in medium with fetal bovine serum and PBS before being injected in 100 µL PBS subcutaneously into the flank of immunodeficient SCID/beige mice (C.B-17/IcrHsd-scid-bg). The mice were bred in the central facility for animal experimentation at the University Medical Center Göttingen under specific pathogen-free conditions in individually ventilated cages and in a 12 hours light-dark cycle. The animal experiments had been approved by the local government and were carried out in compliance with EU legislation (Directive 2010/63/EU). Tumor growth was monitored by palpation and size was recorded using linear calipers. Animals were sacrificed after three months or when a tumour volume of 1 cm^3^ was reached. The tumour volume was calculated by the formula V = πabc/2, where a, b, c are the orthogonal diameters. Autopsies of all animals were performed and half of the tumors were immediately frozen in liquid nitrogen for RNA extraction (Trizol, Thermofisher) followed by qPCR analysis (Applied biosystems, ViiA7). The other half was placed in phosphate-buffered formalin (30 mM NaH_2_PO_4_, 40 mM Na_2_HPO_4_, 4% formalin) for 16 hours before being embedded in paraffin. For histological examination, tissue sections (5 µm) were stained with hematoxylin and eosin (HE) before the slides were scanned with a 20x objective (UPlanApo, NA 0.75) using a dotSlide SL slide scanner (Olympus, Hamburg, Germany) equipped with a peltier-cooled XC10 camera.

### Mass spectrometry analysis

#### Digestion by Filter-Aided Sample Preparation (FASP)

Samples were lysed/denaturated reduced and alkylated during 5 min at 95°C in 100mM Tris/HCl pH8.5, 2% SDS, 10mM TCEP and 50 mM chloroacetamide. For immunoprecipitation, whole samples were digested. For the other samples, protein preparation quality homogeneity and concentration was determined using SDS PAGE separation of an aliquot, colloidal coomassie staining and imaging and integration using Imagelab software (Biorad). 50 µg of proteins were digested as previously described [84]. Briefly, samples were diluted with 8M Urea, 50mM Tris/HCl pH 8.5, transferred onto 30 kDa centrifugal filters and prepared for FASP digestion. Proteins were digested overnight at 37°C with 1µg trypsin (Promega). Peptides were desalted on C18 StageTips, manufactured by stacking six layers of C18 reverse-phase from a disk of 3M Empore Octadecyl C18 High Performance Extraction Disk into a 200 µL micropipet tip.

#### Whole Cell Lysates (WCL) peptide fractionation (BeMI160302 only)

The complex mixture of peptides obtained from FASP were separated in 5 fractions using strong cation exchange (SCX) resin [85]. Briefly, peptides were loaded into pipette-tip columns made by stacking six layers of a 3M Empore cation extraction disk into a 200 µL micropipet tip. Column conditioning was performed using acetonitrile (ACN). We used 0.1% Trifluoroacetic acid (TFA) for column equilibration. Samples acidified with TFA were loaded on the column and washed with 0.1% TFA. Peptides were finally successively eluted using 20% ACN, 0.05% formic acid, ammonium acetate at 75mM, 125mM, 200mM, 300mM. The 5th fraction was eluted in 1.4% NH_4_OH, 80% ACN.

#### Nano Liquid Chromatography-tandem Mass Spectrometry (nLC-MS/MS)

After speed-vaccum drying (Eppendorf concentrator), fractions were solubilized in 10 µL of 0.1% TFA, 10% ACN. Liquid chromatography and mass spectrometry analyses were performed on an U3000 RSLC nanoflow-HPLC system coupled to a Q-Exactive plus Orbitrap mass spectrometer (all from Thermo Fisher Scientific). 1µL of each sample were loaded, concentrated and washed on a C18 reverse-phase precolumn (3µm particle size, 100 Å pore size, 75 µm inner diameter, 2 cm length, Thermo Fischer Scientific) in 2% ACN, 0.1% TFA, 98% H20 at 5 µL/min. After 3 minutes, peptides were separated using a C18 reverse-phase analytical column (2 µm particle size, 100 Å pore size, 75 µm inner diameter, 25 cm length from Thermo Fischer Scientific). A 3 hours and 400 nL/min gradient starting from 99% of solvent A (0.1% formic acid) to 55% of solvent B (80% ACN and 0.085% formic acid) was used for the analysis of samples transfected with shRNAs. A 1 hour gradient starting from 99% of solvent A to 40% of solvent B (80% ACN and 0.085% formic acid) was used for the immunoprecipitation analysis.

The mass spectrometer acquired data throughout the elution process and operated in a data-dependent scheme with full MS scans acquired, followed by up to 10 successive MS/MS HCD-fragmentations on the most abundant ions detected. Spectra were recorded in profile mode. Settings for the immunoprecipitation analysis were: full MS AGC target 1.106 with 60ms maximum ion injection time (MIIT) and resolution of 70 000. The MS scans spanned from 350 to 1500 Th. Precursor selection window was set at 2 Th. HCD Normalized Collision Energy (NCE) was set at 27% and MS/MS scan resolution was set at 17 500 with AGC target 1.105 within 60ms MIIT. Settings for the shRNA analysis were: full MS AGC target 3.106 with 100ms MIIT and resolution of 70 000. The MS scans spanned from 200 to 2000 Th. Precursor selection window was set at 4 Th. HCD Normalized Collision Energy (NCE) was set at 30% and MS/MS scan resolution was set at 17 500 with AGC target 1.105 within 100ms MIIT.

#### Mascot/MyProMS

For IP samples, the peak list of each individual MS/MS spectrum was extracted using Proteome Discoverer 1.4 (Thermo). Identifications (protein hits) were performed by comparison of experimental peak lists with the SwissProt 2017_07 Mus species database using Mascot version 2.5.1. The enzyme specificity was trypsin’s. The precursor mass tolerance was set to 4 ppm and the MS/MS mass tolerance to 20mmu. Cysteins carbamidomethylation was set as constant modification while methionine oxidation was set as variable modification. The lists of protein hits were imported on the MyPROMs software to be compared [86].

#### Maxquant

The mass spectrometry data were analyzed using Maxquant versions 1.5.2.8 and 1.6.1.0 respectively for shRNA and IP analysis [87]. The database used was a concatenation of mouse sequences from the Uniprot-Swissprot database (Uniprot, releases 2016-02, 2017-05 respectively for shRNA and IP analysis) and a list of common contaminants. The enzyme specificity was trypsin’s. The precursor mass tolerance was set to 4.5 ppm and the fragment mass tolerance to 20ppm. Carbamidomethylation of cysteins was set as constant modification and oxidation of methionines as variable modification for all experiments and acetylation of protein N-terminus for IP experiments. Second peptide search was allowed and minimal length of peptides was set at 7 amino acids. False discovery rate (FDR) was kept below 1% on both peptides and proteins. Label-free protein quantification (LFQ) was done using both unique and razor peptides. At least 2 ratio counts were required for LFQ. The statistical analysis was performed using Perseus version 1.6.0.7 [88]. Reverse database hits and contaminants proteins were removed, as well as site modification-only identified proteins. Remaining proteins providing at least two valid values out of three in at least one comparison group were kept for further testing after imputation performed on the missing values. Data imputation step using a random value comprised in the lowest range of LFQ intensities obtained in MaxQuant was performed with the following settings: 0.3 as gaussian width relative to the standard deviation of measured values, and 1.8 as downshift factor (default Perseus values). A non paired t-test was performed to identify differential proteins in shRNA transfected ESC. For identification of USP9X binding partners in co-IPs, unique proteins were identified by MyProMs software and we then calculate a score U/G × (U-G)/(U+G) where U is the number of peptides in USP9X IP and G the number of peptides in the rabbit IgG control. Proteins considered as interacting partners were selected with a score > 1.

The mass spectrometry proteomics data have been deposited to the ProteomeXchange Consortium via the PRIDE partner repository [89].

### Analysis of lactate, pyruvate and nucleotide levels

#### Determination of nucleoside levels

Cellular pellets were deproteinized with an equal volume of 6% perchloric acid (PCA), vortex-mixed for 20 s, ice-bathed for 10 min, and vortex-mixed again for 20 s. Acid cell extracts were centrifuged at 13,000 rpm for 10 min at 4°C. The resulting supernatants were supplemented with an equal volume of bi-distilled water, vortex-mixed for 60 s, and neutralized by addition of 2 M Na_2_CO_3_. Extracts were injected onto a C18 Supelco 5 µm (250 × 4.6 mm) column (Sigma) at 45°C. The mobile phase was delivered at a flow-rate of 1 ml/min using the following stepwise gradient elution program: A–B (60:40) at 0 min→(40:60) at 30 min→(40:60) at 60 min. Buffer A contained 10 mM tetrabutylammonium hydroxide, 10 mM KH_2_PO_4_ and 0.25% MeOH, and was adjusted to pH 6.9 with 1 M HCl. Buffer B consisted of 5.6 mM tetrabutylammonium hydroxide, 50 mM KH_2_PO_4_ and 30% MeOH, and was neutralized to pH 7.0 with 1 M NaOH. Detection was done with a diode array detector (PDA). The LC Solution workstation chromatography manager was used to pilot the HPLC instrument and to process the data. Products were monitored spectrophotometrically at 254 nm, and quantified by integration of the peak absorbance area, employing a calibration curve established with various known nucleosides. Finally, a correction coefficient was applied to correct raw data for minor differences in the total number of cells determined in each culture condition.

#### Determination of Lactate and Pyruvate levels

Collected cell culture media were deproteinized with ice-cold 4% (w/v) perchloric acid and after centrifugation, the metabolites were assayed on neutralized perchloric filtrates by enzymatic methods and spectrophotometry according to previous protocols [90]. Briefly, lactate was converted into pyruvate with NAD and LDH in a hydrazine buffer and the resulting NADH is measured by spectrophotometry. Pyruvate was also measured by exploiting the reverse reaction with NADH and LDH and in which NADH will be oxidized into NAD. NADH consumption is assayed by spectrophotometry. Data are reported as mole per cell; and the ratio lactate / pyruvate has been calculated for each sample.

#### Determination of succinate ubiquinone oxido-reductase, cytochrome C oxidase and citrate synthase activities

Enzymatic activities were determined using conventional protocols previously described [91]. Enzymatic activities were normalized to the amount of proteins in the lysate and are expressed as nanomoles/min/mg of proteins.

### Mitochondiral respiration analysis

Oroboros and Seahorse analysis of OXPHOS and glycolysis were conducted as previously described [66]. Data were normalized to cell number in each experiment.

### qPCR quantification of mitochondrial genome abundance

RNA and DNA from frozen Usp9X^+/y^ and Usp9X^E2/Y^ ESC pellets were obtained by following the protocol from the Qiagen RNA/DNA extraction kit (Qiagen, 80204). DNA was diluted to 100 pg for qPCR analysis on a LightCycler® 96 System available on the Genom’ic facility at Institut Cochin.

Primers for DNA repeated sequences (LINE, LTR, SINE) and the mitochondrial genome are available in Supplementary Table 7. qPCR values were corrected for primer efficiency. Ct values for Usp9X^+/y^ and Usp9X^E2/Y^ samples were normalised to Ct values abtained from a third murine ESC cell line grown at the same time.

### Microarray analysis

Purified RNAs from V6.5 ESC electroporated with shRNA vectors directed against Usp9X or control sequences (i.e. Icos or linker) were processed by the GENOM’IC platform at Institut Cochin. Three independent biological samples from each group were processed and hybridized to the Affymetrix WT plus MouseGene2.0 array. After validation of the RNA quality with Bioanalyzer 2100 (using Agilent RNA6000 nano chip kit), 250 ng of total RNA were reverse transcribed following the GeneChip® WT Plus Reagent Kit (Affymetrix). Briefly, the resulting double strand cDNA were used for *in vitro* transcription with T7 RNA polymerase (all these steps are included in the WT cDNA synthesis and amplification kit of Affymetrix). After purification according to Affymetrix protocol, 5.5 μg of Sense Target DNA were fragmented and biotin labelled. After control of fragmentation using Bioanalyzer 2100, cDNA was then hybridized to GeneChip^®^ Mouse Gene 2.0 ST (Affymetrix) at 45°C for 17 hours. After overnight hybridization, chips were washed on the fluidic station FS450 following specific protocols (Affymetrix) and scanned using the GCS3000 7G.

The scanned images were then analysed with Expression Console software (Affymetrix) to obtain raw data (cel files) and metrics for Quality Controls. Raw data were normalized using the Robust Multichip Algorithm (RMA) in Bioconductor using Brain array EntrezGene CDF version 21 for annotations. Then all quality controls and statistics were performed using Partek Genomic Suite. To find differentially expressed genes, we applied a one-way analysis of variance (ANOVA) for each gene. Then, we used *p*-values and fold changes to filter and select differentially expressed genes.

### Immunoprecipitation

ESC (maintained in LIF + serum; treated with MG132 or MG132+WP1130; at Day 1 and 3 post-LIF removal) were lysed in IPH buffer (50mM Tris HCl pH8, 150mM NaCl, 5mM EDTA pH8, 0,5% NP-40) with 30U of benzonase (E1014-5KU, Sigma, or 70664-10KU, Novagen) for 1 hour on ice and then pelleted at 13000rpm, 10min, 4°C. Supernantants were used for immunoprecipitation over-night with relevant antibodies or irrelevant IgG bound to Dynabeads Protein G magnetic beads (ThermoFisher Scientific, 10004D). Washes were done 6 times with 1 mL of cold IPH buffer. Elution is performed in pre-warmed 1% SDS 15 min in thermomixer at 850rpm. Samples (input and immunoprecipitates) were then analyzed by SDS-Page followed by western blotting or by PageBlue^TM^ protein staining (ThermoFisher Scientific, 24620) or by mass-spectrometry.

### Comparison between Usp9X target genes and ChIP-sequencing datasets

ChIP-sequencing data for ESRRB, NANOG, SOX2 and SUZ12 were retrieved from NCBI GEO GSE11431 [69]. Data were processed for peak calling with MACS2 using default parameters and input as control [92]. Annotation of the binding sites was done using the R/Bioconductor package ChIPseeker [93]. We then intersected intervals from ChIP-sequencing data and Usp9X target genes using the mus musculus genome GRCm38/mm10 (release December 2011) as reference.

### Annotation of datasets: GO terms, compartments and mammalian phenotypes annotation

The mammalian phenotype (MP term), anatomical (EMAPA term), molecular (GO MP) and cellular (GO BP) function annotation of USP9X potential interactors was performed using the Mouse Mine interface (http://www.informatics.jax.org/) with Holm-Bonferroni adjustment [94]. GO term enrichment, pathway analysis and comparison with chemical and genetic gene-set signatures were performed using the GSEA and molecular signatures database or MSigDB collection (http://software.broadinstitute.org/gsea/msigdb/index.jsp) with *P*-value < 1e-5 and FDR < 0.01 [95]. StemChecker was utilized to annotate 25 stemness signatures and 73 TF ChIP sets [96]. Among the 311 USP9X transcriptional target genes, 65 were not found in the database and 36 were found but not in any of the selected datasets. Ingenuity Pathway Analysis (QIAGEN Inc.) and PantherDB (version 14.1 released 2019-03-12) with Fisher exact-t-test and FDR were utilized to functionally annotate transcriptome and proteome from Usp9X^KD^ ESC [97, 98]. Protein subcellular annotation was performed using the manually curated “COMPARTMENTS” web interface [99].

### Transcription factor motif search

Motif enrichment analysis were performed using the oPOSSUM3 interface based on Ensembl release 64 (Sep 2011) [100]. We utilized the following criteria: conservation 0.4, matrix score threshold: 85%, upstream/downstream: 2000 base pairs, z-score ≥ 10 and Fisher score ≤ 7 with a minimum score of 8-bits. The 29347 mouse genes in the oPOSSUM database were used as background for the analysis.

### Expression profile of ESC in “2i”, LIF+Serum, in differentiation and following transcription factor depletion

Expression data from J1 WT mouse ESC cultivated in serum medium and after two weeks in 2i medium in presence of vitamin C were obtained from the NCBI GeneOmnibus repository, GSE71591 [62]. Expression data from ESC and spontaneously differentiated ESC were kindly shared by Dr Matthieu Gerard and freely re-usable (Institut de Biologie Intégrative de la Cellule, Commissariat à l’Energie Atomique et aux Energies alternatives, France). Expression data from wild-type murine ESC at day 0 and day 8 of differentiation into embryonic bodies were obtained from GSE96809 [59]. Expression data from wild-type D3 ESC at day 0 and day 8 of differentiation into embryonic bodies were obtained from GSE124241 [60]. Expression data from E14 WT ESC derived and maintained in serum or 2i were downloaded from SRR064970 and SRR064972 respectively [61]. Expression data from ESC [MC1R(20)] transfected with shRNAs against Esrrb, Sall4, Sox2 and Pou5f1 were downloaded from NCBI GEO GSE26520 [9].

The expression difference is considered significant for a fold change of ≤-1.2 or ≥1.2. Data across different publications were combined based on matching their Entrez Ensembl identifiers. Gene name conversion was performed using g:Profiler [101].

### Data availability

The gene expression data produced in this study are available at the NCBI Gene Expression Omnibus. The mass spectrometry proteomics data have been deposited to the ProteomeXchange Consortium via the PRIDE partner repository.

## Acknowledgements

We thank the CYBIO, GENOM’IC and 3P5 facilities at Institut Cochin (Paris, France) for training and assistance with cytometry, transcriptomics and proteomics analyses. BM and MdD thank Catherine Gayed, Benjamin Martin and Nyota Kunyu for performing experiments during their short-term internships in the laboratory. We are grateful to Gwenneg Kerdivel, Lélia Polit (Institut Cochin, France) and Olivier Kirsh (UMR7216, France) for bio-informatical help and Pablo Navarro (Institut Pasteur, Paris, France), Didier Trouche (Centre de Biologie Intégrative, Toulouse, France) and Constance Ciaudo (EH Zurich, Switzerland) for insightful discussions and suggestions during the conduct of this project.

## Supplementary Figures

**Supplementary Figure 1:**
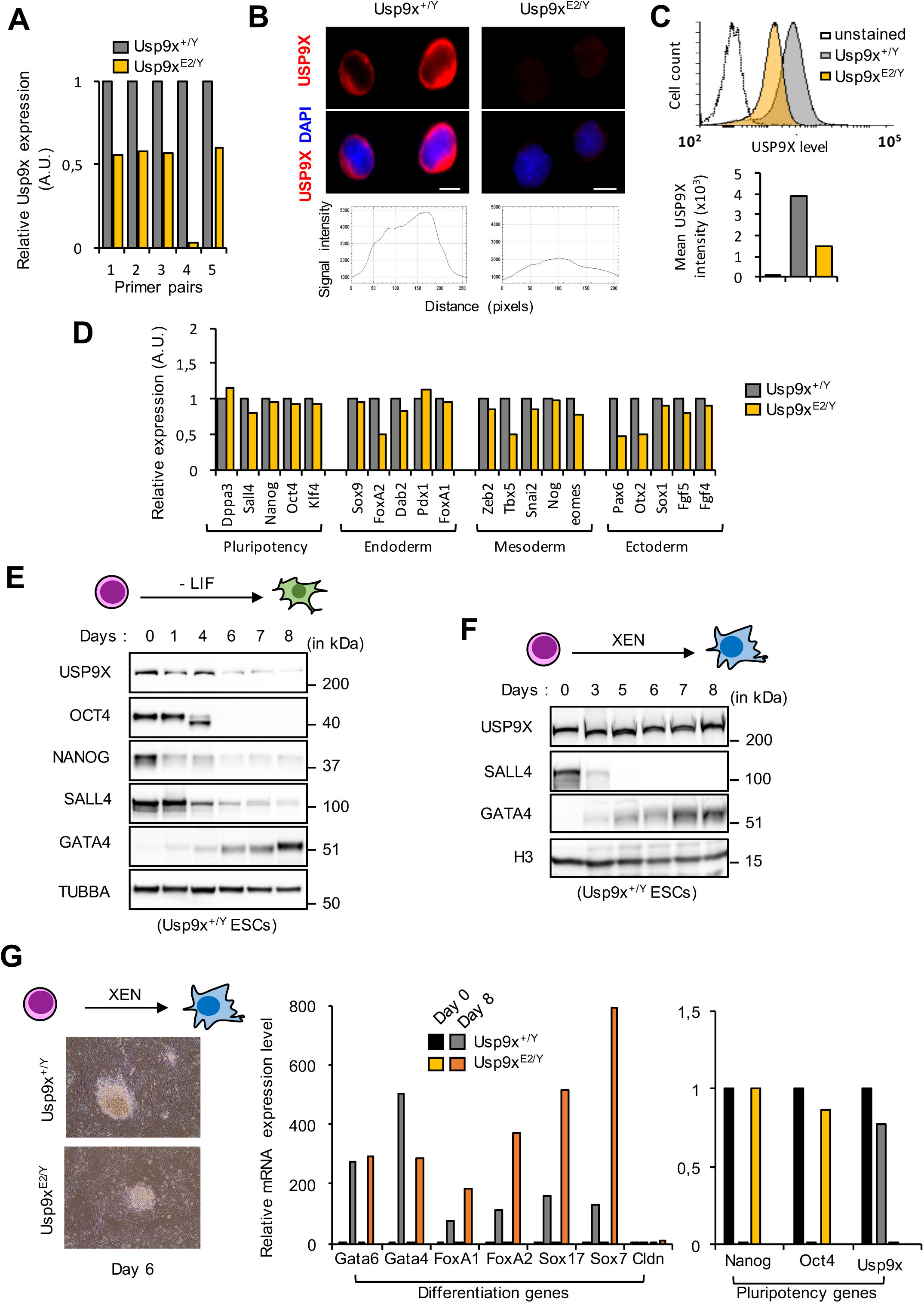
Characterization of USP9X^E2/Y^ ESC. (**A**) qRT-PCR analysis of Usp9X expression in USP9X^E2/Y^ and control ESC cultured in serum + LIF media (n = 3). Usp9X expression is normalized to GAPDH expression with value set up to 1 in USP9X^+/y^ ESC. Primer pairs are as follow: 1, overlapping exon junction 5-6; 2, overlapping exon junction 9-10; 3, overlapping exon junction 17-18; 4, overlapping exon junction 2-3; and 5, overlapping exon junction 3-4. (**B**) Immunofluorescence analysis of USP9X in USP9X^E2/Y^ and control ESC. USP9X was stained by indirect immunofluorescence using an Alexa-FLUOR Plus 555-secondary antibody (in red). DNA is labelled with DAPI (in blue). Relative quantification of USP9X signal intensity is provided (i.e. the x-axis represents the horizontal distance through the images and the y-axis the vertically averaged pixel intensity). Scale bar: 2 μm. (**C**) Flow cytometry analysis of USP9X levels in USP9X^E2/Y^ and control ESC cultured in serum + LIF media. Representative experiment and USP9X amount are shown. (**D**) RT-PCR analysis of pluripotency (Dppa3, Sall4, Nanog, Oct4 and Klf4), endoderm (Sox9, FoxA2, Dab2, Pdx1 and FoxA1), mesoderm (Zeb2, Tbx5, Snai2, Nog and Eomes) and ectoderm (Pax6, Otx2, Sox1, Fgf5 and Fgf4) markers in Usp9X^E2/Y^ and control ESC maintained in LIF + serum medium. (**E**) Kinetics of USP9X, OCT4, NANOG, SALL4, GATA4 and TUBBA abundance during differentiation of USP9X^+/y^ ESC into embryonic bodies. Differentiation was induced by removal of LIF at day 0 and addition of retinoic acid (5 µM) at day 4 and 6. As anticipated, pluripotency factors OCT4, NANOG and SALL4 are downregulated during differentiation and GATA4 abundance rises starting on day 4. (**F**) Kinetics of USP9X abundance during differentiation of USP9X^+/y^ ESC into XEN cells. As anticipated, pluripotency factors SALL4 is downregulated during differentiation and GATA4 abundance rises starting on day 3. (**G**) Representative extra-embryonic cells (XEN) derived from USP9X^E2/Y^ and control ESC seen at day 6 of differentiation. RT-PCR analysis of pluripotency (Oct4 and Klf4) and differentiation (Gata6, Gata4, FoxA1, FoxA2, Sox17, Sox7 and Cldn) markers in USP9X^E2/Y^ and control ESC at different days after induced differentiation (same protocol as in panel 1F).

**Supplementary Figure 2:**
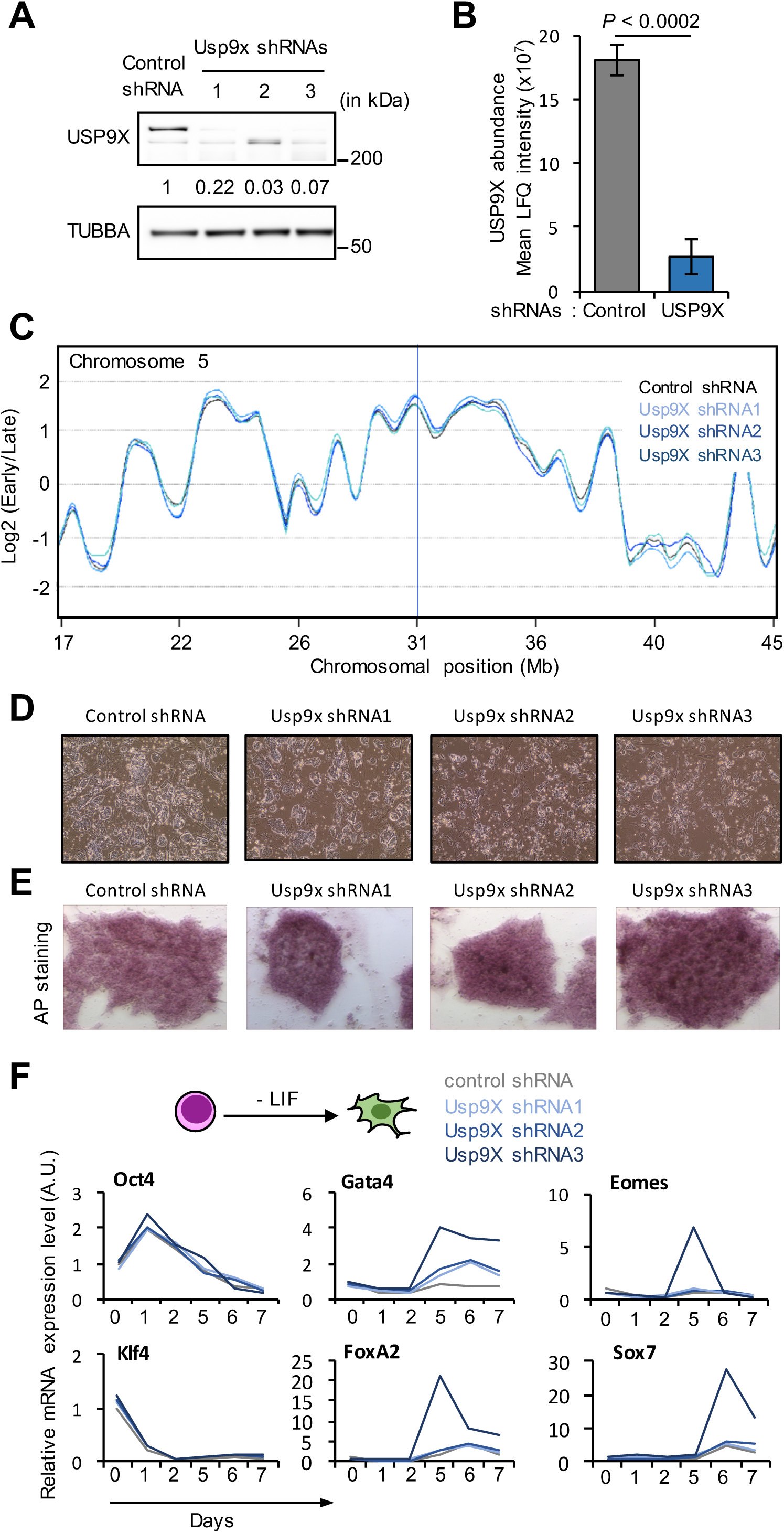
Characterization of USP9X^KD^ ESC. (**A**) Western blot analysis of USP9X and TUBBA levels in ESC transfected with 3 independent shRNA constructs targeting Usp9X or a control shRNA vector. Relative amount of USP9X protein normalised to TUBBA is indicated. (**B**) Label-free quantification (LFQ) of USP9X protein levels in ESC transfected with control shRNA or Usp9X shRNAs. Data are presented as mean ± SD (n = 3). *P*-value from t-test vs control. (**C**) Example of DNA replication timing profiles in Usp9X^KD^ and control ESC. Probe values represented as log2(early/late) along murine chromosome 5 revealed no significant difference between the two cell types. Mb, mega-bases. (**D**) Representative images of ESC after transfection of control shRNA or Usp9X shRNAs. Subtle morphological changes are detectable at the periphery of the ESC colonies. (**E**) Representative alcaline phosphatase (AP) staining of ESC after transfection of control shRNA or Usp9X shRNAs. (**F**) qRT-PCR analysis of pluripotency (Oct4 and Klf4) and differentiation (FoxA2, Gata4, Sox7 and Eomes) markers in ESC transfected with control shRNA or Usp9X shRNAs and further induced to differentiate into embryonic bodies by LIF removal.

**Supplementary Figure 3:**
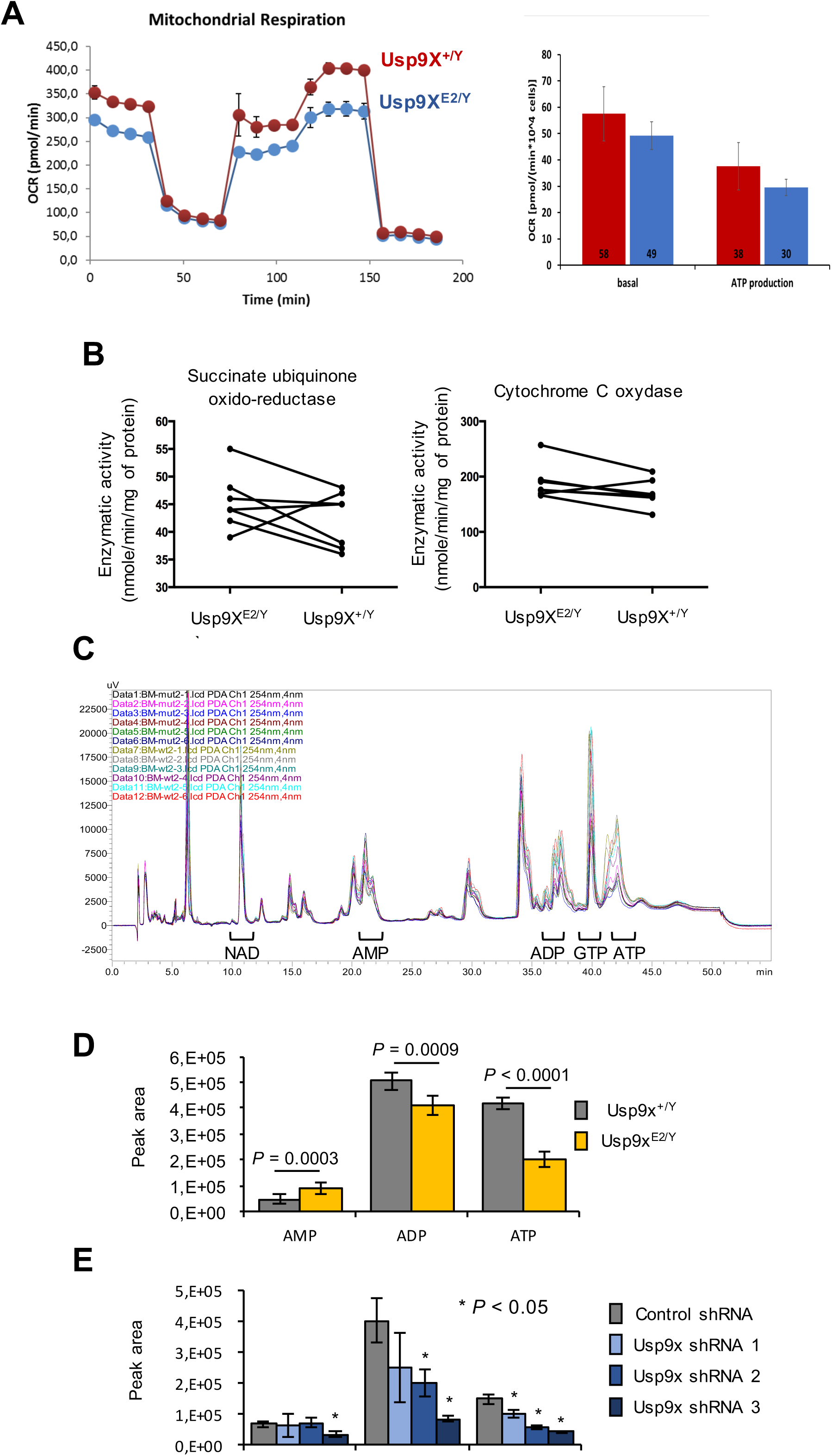
Depletion of Usp9X impairs ATP levels in ESC. (**A**) Analysis of mitochondrial activity by Seahorse. Left Panel: representative measurements. Right panel: Graph reporting basal cell respiration, ATP-production respiration, leak and respiratory reserve of Usp9X^E2/Y^ and control ESC (n = . (**B**) Measure of succinate ubiquinone oxido-reducate (left graph) and cytochrome C oxidase (right panel) activities in Usp9X^E2/Y^ and control ESC (n = 6). Results were not statistically significant between groups (paired t-test). (**C**) HPLC data used for the measure of AMP, ATP and ADP levels in Usp9X^E2/Y^ and control ESC. (**D**) Quantification of AMP, ATP and ADP levels in Usp9X^E2/Y^ and control ESC. Area under the curve was determined from HPLC data and represented as mean (±SD) values (n = 6). (*p*-value from t-test, Usp9X^E2/Y^ vs Usp9X^+/y^). (**E**) Quantification of AMP, ATP and ADP levels in ESC transfected with Usp9X and control shRNAs. Data are represented as mean (±SD) values (n = 3).

**Supplementary Figure 4:**
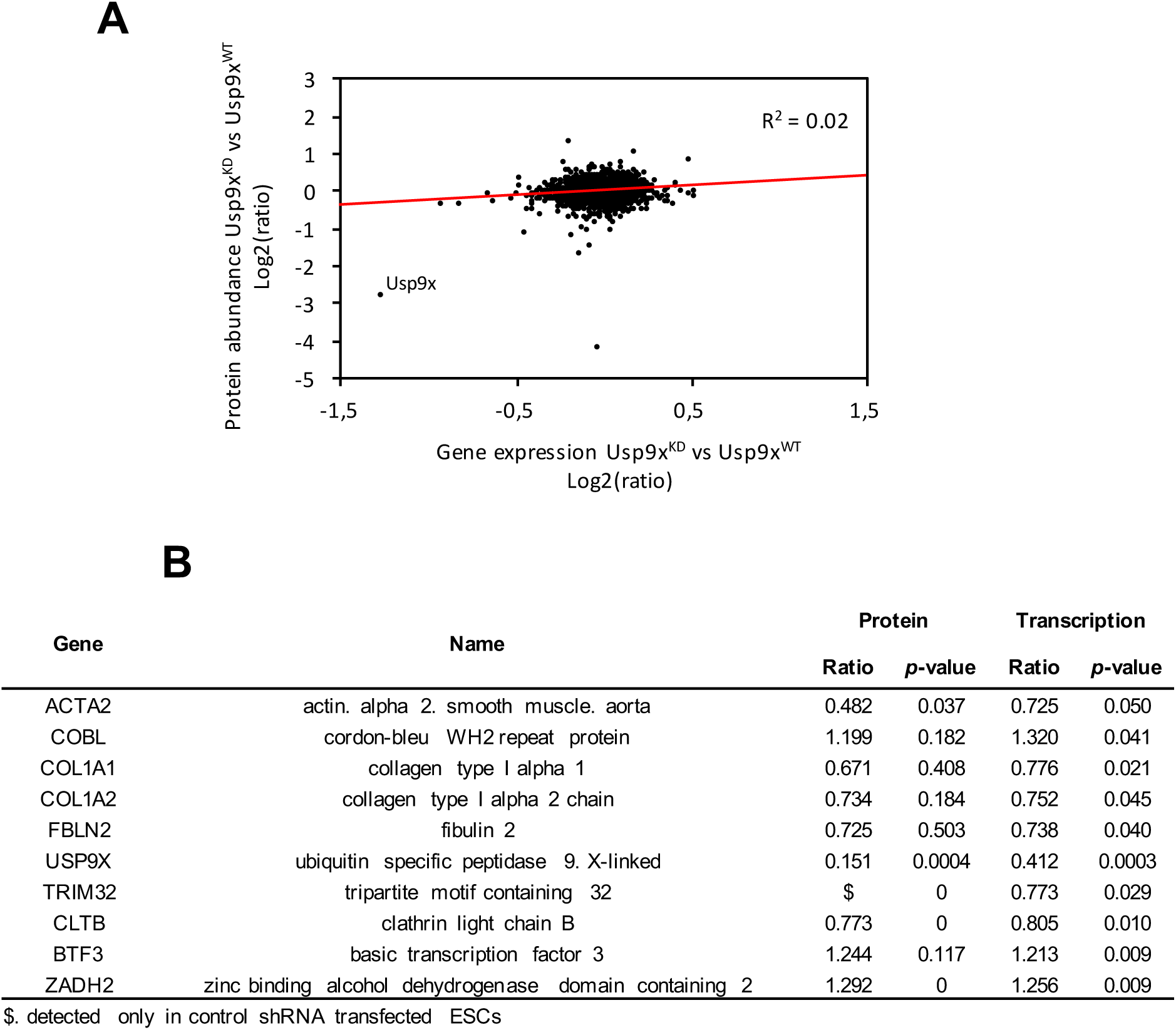
Most proteins regulated by USP9X are not transcriptional target. (**A**) Correlation between protein and mRNA fold change in Usp9X^KD^ ESC and control ESC. (**B**) List of the 10 proteins showing correlation between mRNA and protein regulation upon Usp9X depletion.

**Supplementary Figure 5:**
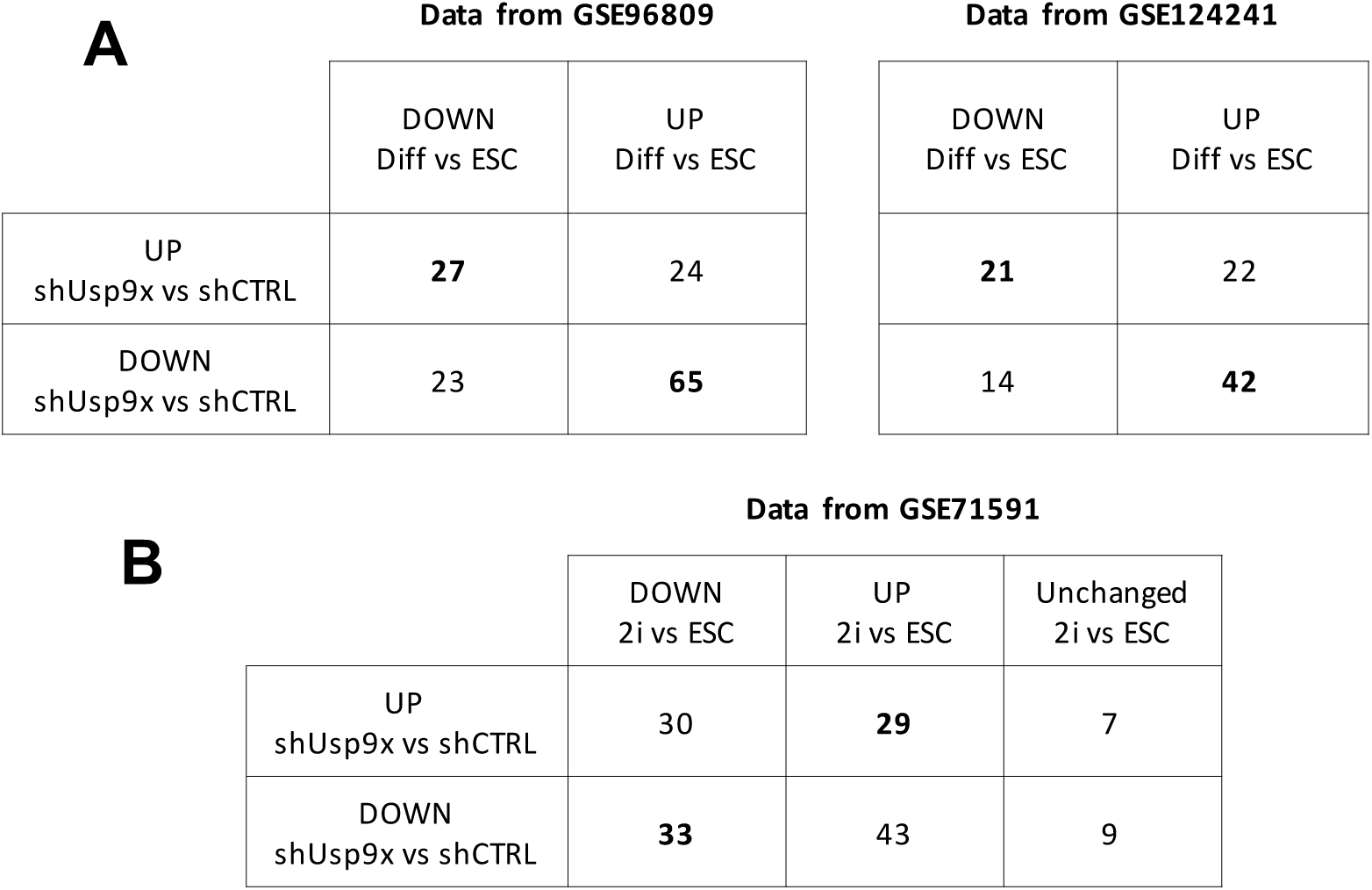
Characterization of USP9X^KD^ transcriptome. (**A**) Intersection of differentially expressed genes identified from USP9X^KD^ ESC (vs control ESC) and micro-array datasets of differentially expressed genes during ESC differentiation from GSE96809 and GSE124241. (**B**) Intersection of differentially expressed genes identified from USP9X^KD^ ESC (vs control ESC) and a micro-array dataset of differentially expressed genes during LIF + serum to 2i transition reported in GSE71591.

**Supplementary Figure 6:**
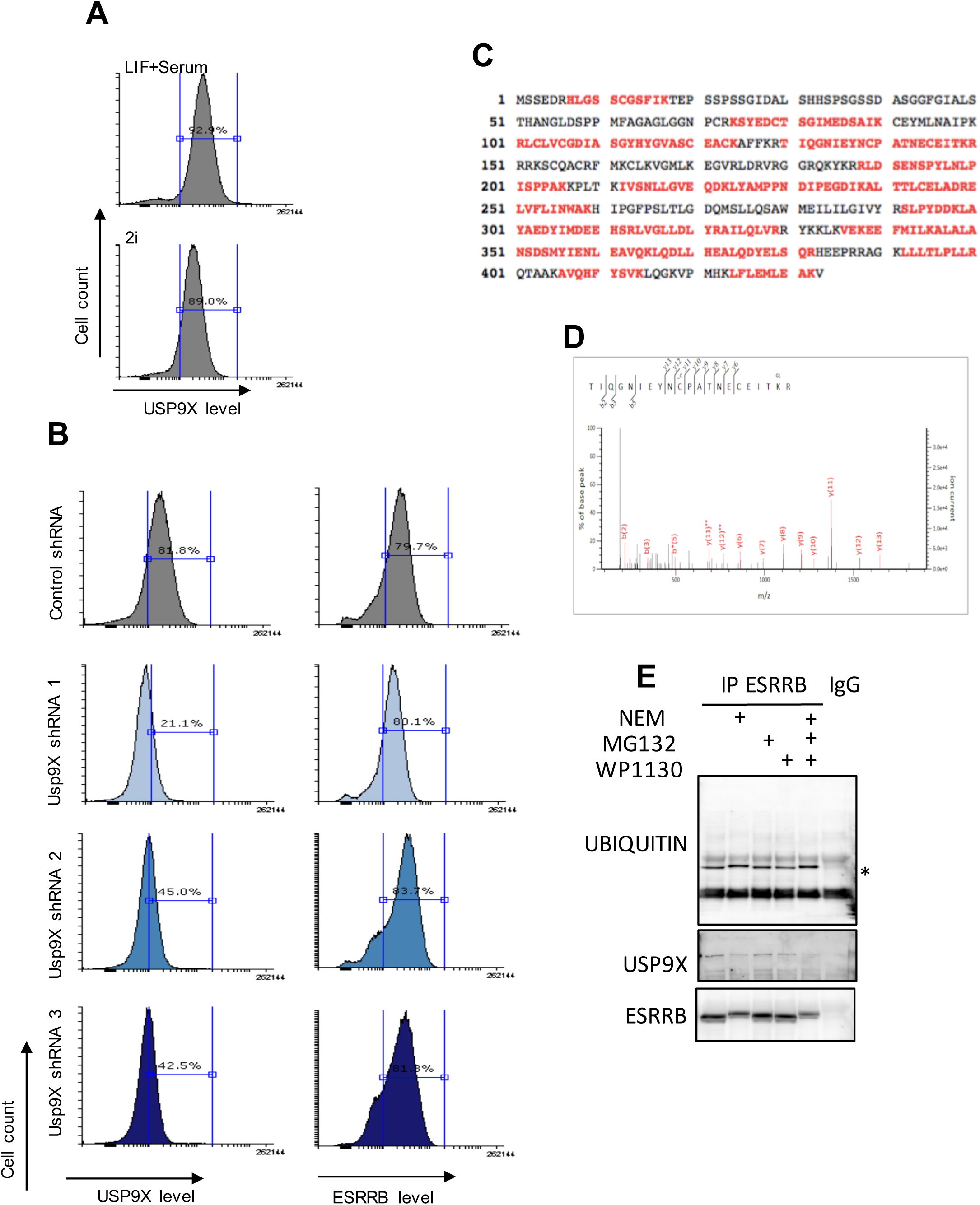
ESRRB is a ubiquitinated protein. (**A**) Sequence coverage of ESRRB from a representative mass spectrometry run. Detected peptides are highlighted in red. Sequence coverage is similar whether ESRRB is immunoprecipitated from ESC treated with MG132 (10 mM for 4 hours) or not. (**B**) MS/MS spectrum of the ubiquitinated peptide detected in our mass-spectrometry analysis, in samples from ESC treated with MG132 (10 mM for 4 hours) or not. (**C**) V6.5 ESC were treated with MG132 at 10 mM for 4 hours or WP1130 at 5 μM for 4 hours prior to cell lysate, or with NEM during lysate preparation, and ESRRB immunoprecipitation. Ubiquitination level of ESRRB was then assessed by western blotting using an anti-ubiquitin antibody. (**D**) Flow cytometry analysis of USP9X levels in ESC cultured in LIF + serum (top panel) or 2i (bottom panel). Hundred thousand cells were analysed per condition. (**E**) Flow cytometry analysis of USP9X (left panels) and ESRRB (right panels) levels in V6.5 ESC transfected with Usp9X or control shRNAs. Hundred thousand cells were analysed per condition.

